# Solvent-Isotope Effects in Biomolecular Phase Separation and Fibrillation of Disordered Proteins and Peptides

**DOI:** 10.64898/2026.09.08.748679

**Authors:** Hanuman Singh, Zayne Yeager, Morgane Herlory, Sayanta Mahapatra, Maciej A. Walczak

**Affiliations:** Department of Chemistry, University of Colorado, Boulder, CO 80309, United States

## Abstract

Heavy water (D_2_O) is widely used in biomolecular spectroscopy and imaging, often under the assumption that it is an inert replacement for H_2_O. However, D_2_O differs subtly in hydrogen-bonding, viscosity, and dielectric properties, which can alter biomolecular interactions and self-assembly. Here, we test how solvent isotope substitution modulates protein/peptide phase separation and amyloid formation in multiple intrinsically disordered systems. Using turbidity-based phase diagrams and microscopy, we quantify how D_2_O shifts protein-RNA complex coacervation boundaries and alters condensate morphology. Droplet recovery measurements indicate significant solvent-dependent changes in condensate material properties. We further evaluate amyloid formation kinetics, in the presence or absence of a cofactor, supported by orthogonal structural characterization, and assess the functional consequences of tau fibrils using a tau biosensor seeding assay with explicitly defined seed delivery conditions. Together, these results show that D_2_O can systematically bias liquid-liquid phase separation and aggregation readouts and should be treated as an active experimental variable rather than a neutral solvent substitute.

## INTRODUCTION

Liquid-liquid phase separation (LLPS) has emerged as a central biophysical principle that explains how cells compartmentalize reactions, buffer stress and choreograph signaling without the need for bounding membranes.^1-3^ In vitro reconstitution and quantitative imaging have revealed that a strikingly wide spectrum of macromolecules such as intrinsically disordered proteins (IDPs),^4, 5^ modular multivalent proteins,^6^ RNA,^7^ and small metabolites^8^ can demix from the surrounding cytoplasm or nucleoplasm to form dynamic, fluid condensates whose material properties are sensitive to weak non-covalent interactions.^7^ As the catalogue of phase-separating scaffolds has grown, so too has the range of analytical tools deployed to study them.^9-11^ Heavy water (D_2_O) is commonly used in biomolecular spectroscopy, imaging, and scattering to improve contrast, reduce background, or simplify spectral analysis.^12^ Although often treated as a near-equivalent replacement for H_2_O, D_2_O differs in hydrogen-bond dynamics, viscosity, dielectric properties, surface tension, and hydration behavior.^13-15^ Protein-solubility studies have shown that globular proteins can exhibit altered effective interactions, precipitation thresholds, and phase behavior in D_2_O.^16-22^ Recent work has extended this principle to synthetic coacervates.^23^ Together, these observations establish solvent isotope composition as an active physicochemical variable rather than an inert experimental substitution. Beyond their conceptual interest, solvent-isotope effects carry immediate practical implications. Drug-discovery campaigns that target disease-associated condensates,^24^ such as the solidifying assemblies of tau^25^ or the transcriptional hubs that drive oncogene expression,^26, 27^ routinely conduct high-throughput screens in partially deuterated buffers. Heavy water stabilizes or rigidifies droplets, and apparent hits selected under those conditions may fail in protonic buffers or in vivo, where the energetic landscape differs.

The physical origins of the phenomenon are multifaceted and rooted in nuclear-quantum effects.^13, 14, 28^ The larger mass of deuterium lowers the zero-point vibrational energy of the O-D bond, strengthening individual hydrogen bonds propagating through the extended H-bond network of water.^15^ As a consequence, D_2_O forms a more structured and less mobile hydrogen-bond network than H_2_O, which can reduce protein hydration and protein–water hydrogen bonding while increasing the energetic cost of solvent reorganization associated with hydrophobic cavity formation.^13, 29-31^ LLPS, which relies on a delicate competition between solvation free energy in the dilute phase and the multivalent “sticker” interactions in the dense phase, is therefore highly susceptible to even modest isotope-induced shifts in solvent energetics. The greater energetic cost of solvent reorganization in D_2_O increases the tendency of hydrophobic patches to associate, strengthening effective hydrophobic interactions. Because increased protein sticker hydrophobicity often enhances phase separation, and in some systems, drives the formation of solid-like assemblies, the mechanistic consequences of isotopically enhanced hydrophobic interactions in coacervates offer an interesting area for further exploration.^32^ Here, we report chemical, biophysical and cellular studies that closes these gaps and establishes a predictive foundation for solvent-isotope control of biomolecular phase separation and protein fibrillation.

## RESULTS

### Design and preparation of molecular probes

To investigate the effects of solvent isotopes on the structural and biophysical properties of proteins and peptides, we employed a combination of methods, including synthetic chemistry, biolayer interferometry (BLI) to evaluate the binding affinity of different proteoforms to various RNA species, turbidity assays with RNA to monitor LLPS, FRAP to characterize droplet fluidity,^33^ and aggregation assays to assess structural changes. For these studies, we assembled a panel of constructs comprising peptides, glycopeptides, and proteins (Figure 1). As a model intrinsically disordered protein, we used microtubule-associated protein tau in its longest isoform (2N4R tau), as well as shorter fragments that are known to undergo heparin-induced fibrillation (K18 and tau(291-391)), and tau(287-391), which can undergo self-aggregation at high protein concentrations without the need for additional polyanions.^34^ Tau is a model for studying isotope effects on electrostatically driven complex coacervation,^35-37^ and the transition of condensed liquid droplets into amyloids. A model peptide **R1**, (RRASL)_3_, which can form condensates in vitro,^38-40^ was chemically modified by mutating alanine into more aromatic residues such as phenylalanine (**R2**) and tryptophan (**R3**). These modifications allow to delineate the impact of solvent isotope effects on aromatic-cation interactions.^41-46^ Glycosylated peptides **R4**-**R8** were selected as a model to understand the role of (poly)hydroxylated moieties in phase separation. Our recent work demonstrates that *N*-linked glycans can have a profound effect on in vitro condensation, and, in some cases, may even completely suppress condensate formation.^47^ On the other hand, O-linked glycans remain poorly studied although reports with O-GlcNAc proteins such as RNA-binding protein EWS indicate the effects of condensation to be in general suppressive.^48^ Glycosylation influences LLPS through several mechanisms including steric hindrance inducing conformational and strong hydrogen-bond effects.^49, 50^ Attaching sugars increases the overall hydrophilicity and induce local conformational changes by restricting backbone flexibility or altering secondary structures.^51^ Moreover, glycans can engage in hydrogen bonding with the protein backbone or side chains, which might stabilize certain folds or block intramolecular hydrogen bonds that would otherwise promote aggregation.^52-55^ To this end, the model peptide (RRASL)_3_ was modified with one or three α-GalNAc resides (**R4** and **R7**, respectively), and the glycans were introduced into the hydrophobic peptide (**R8**). To understand if the identity of the saccharide can influence the LLPS, we replaced GalNAc with β-D-galactose (**R5**) and β-D-glucose (**R6**). Finally, HBP-1 peptide **R9**, which is derived from histidine-rich squid beak proteins (HBPs), was included because it undergoes self-coacervation through hydrophobic and π-π stacking interactions.^56^

**Figure 1.**
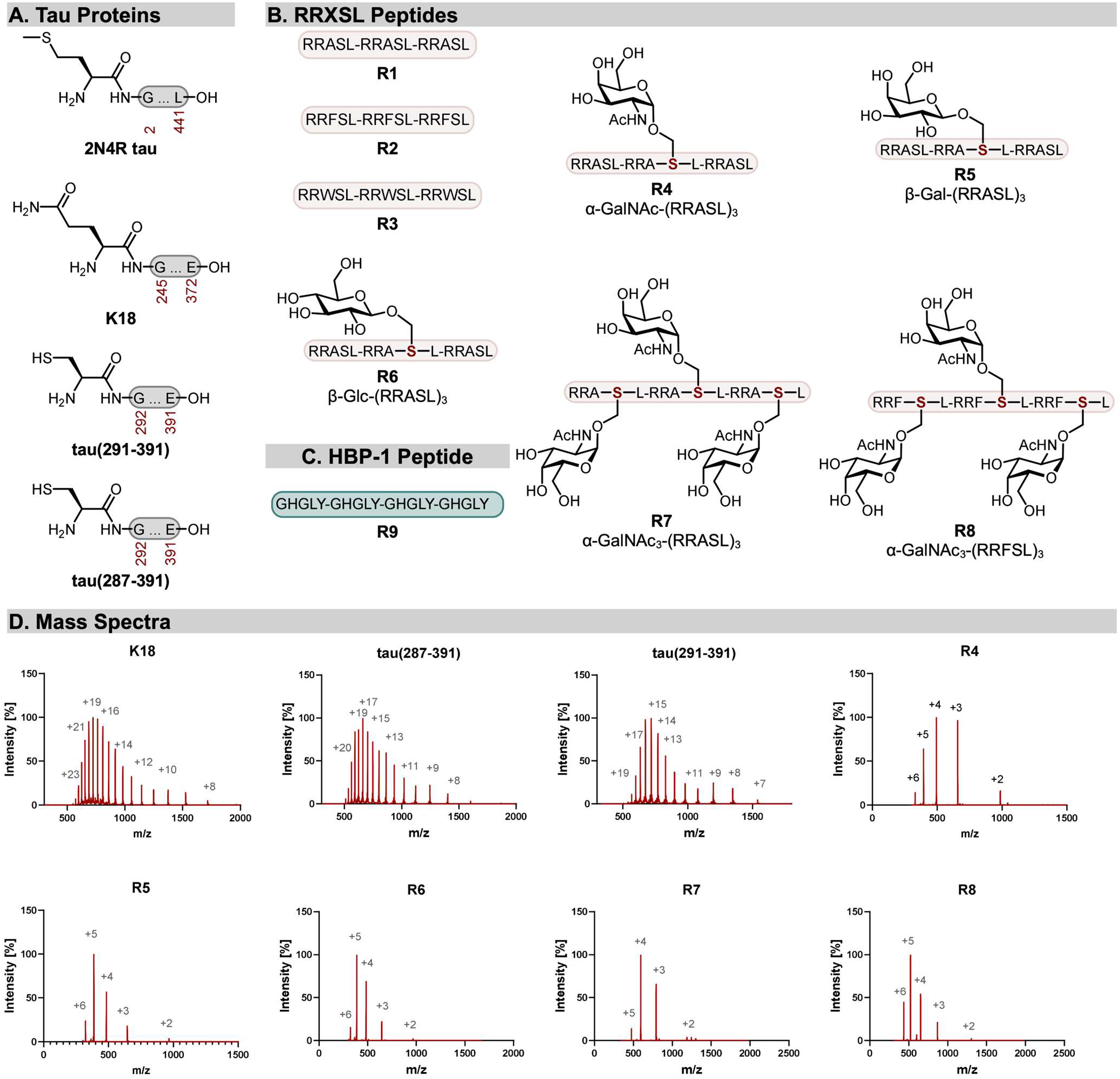
Peptide, glycopeptide, and tau constructs used to evaluate solvent-isotope effects on biomolecular self-assembly. (A) Tau proteins and fragments used in this study: full-length 2N4R tau, K18 tau, tau(291-391), and tau(287-391). (B) RRXSL-derived peptides and glycopeptides R1–R8 designed to test the effects of charge density, aromaticity, glycan identity, and glycan number on peptide/RNA phase separation. (C) HBP-1-derived peptide R9 used as a model of salt-tolerant self-coacervation. (D) Representative MS spectra confirming the identities of synthetic peptides, glycopeptides, and tau fragments.

Proteins 2N4R tau, K18, tau(291-391), and tau(287-391) were prepared according to previously reported procedures,^47, 57^ while a series of RRASL, its sugar-modified variants with an increasing number of monosaccharide molecules were designed and synthesized using solid-phase peptide synthesis (Figure 1B). Prior to all experiments in heavy water, all reagents were repeatedly lyophilized from D_2_O (98% D) to remove all exchangeable protons.

### RNA-induced phase separation

To investigate the ability of RRASL (**R1**) and its variants to undergo phase separation in the presence of RNA, we monitored both turbidity formation and condensate morphology upon addition of poly(U) under matching pH/pD conditions (Figure 2). In the case of **R1**, complex coacervation is driven by attractive electrostatic interactions between positively charged residues in the peptide and polyanionic RNA(Figure 2A). **R1** exhibited a pronounced turbidity increase upon mixing with poly(U), which was accompanied by a visible change in solution appearance, indicative of condensate formation and confirmed the presence of micron-sized, spherical droplets (Figure 2B-D). After centrifugation, the droplet-rich phase settled at the bottom of the tube, further confirming condensate formation. Next, we employed a turbidity phase diagram assay to determine how proteins and peptides undergo polyanion-induced LLPS in H_2_O versus D_2_O (Figure 2E). Turbidity profiles of **R1** revealed pronounced solvent-dependent differences: in D_2_O, the construct exhibited a sharp increase in turbidity followed by saturation, whereas in H_2_O the maximum turbidity was consistently reduced and shifted to higher RNA ratios. These results demonstrate that heavy water enhances the efficiency of condensate formation relative to normal aqueous conditions for electrostatically driven LLPS, likely due to altered hydrogen-bonding and solvation dynamics that modulate peptide-RNA interactions. Fluorescence recovery after photobleaching (FRAP) was performed on **R1** droplets under both solvent conditions (Figure 2F and 2G). In H_2_O, **R1** droplets exhibited rapid and nearly complete fluorescence recovery indicating a highly dynamic, liquid-like internal environment. By contrast, droplets formed in D_2_O displayed markedly slower recovery kinetics and significantly reduced mobile fractions, consistent with a transition toward a more gel-like state. The circular dichroism (CD) spectra of **R1** in the presence of poly(U) revealed that a highly disordered structure in both solvents (Figure 2H).

**Figure 2.**
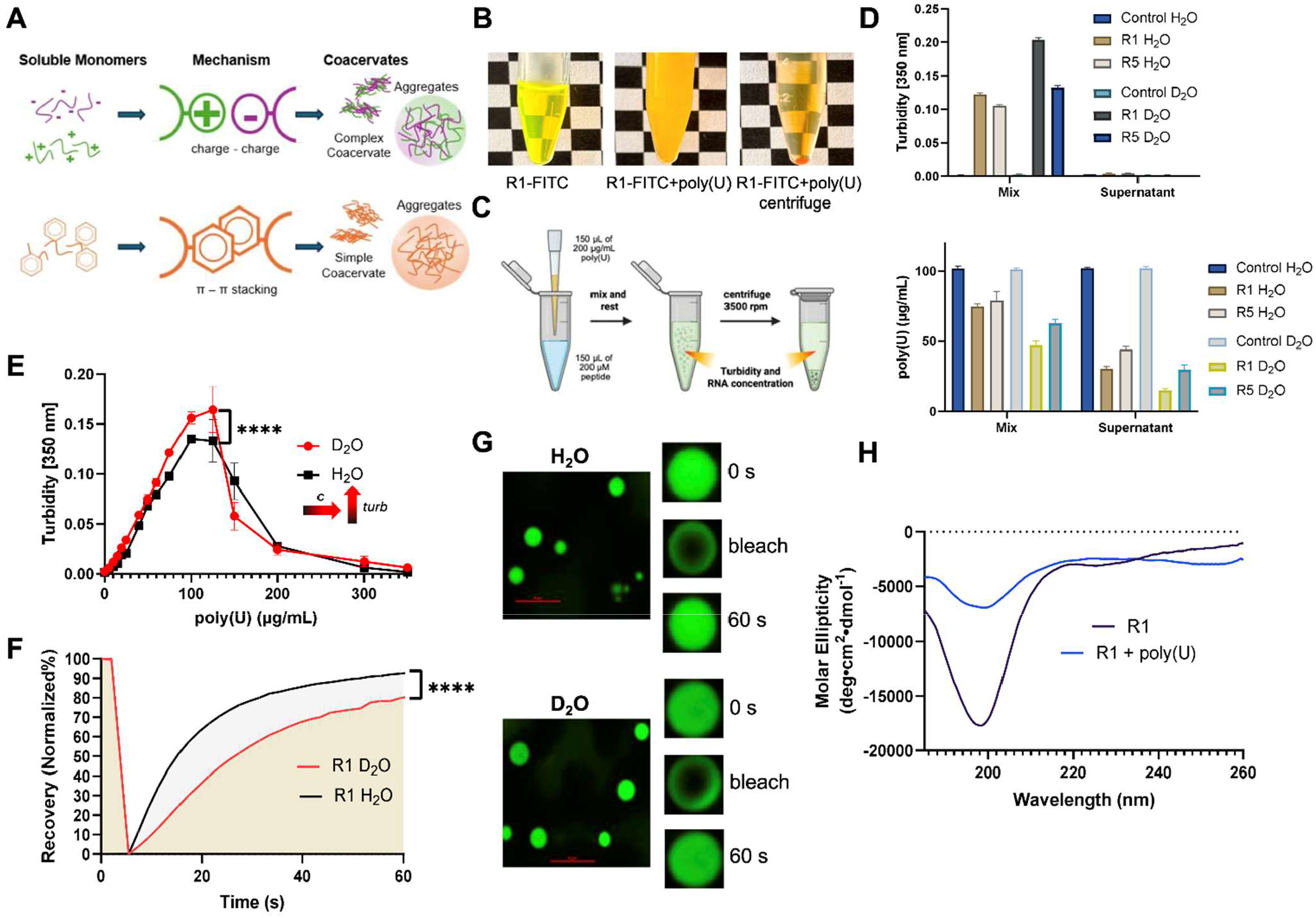
D_2_O enhances R1/poly(U) complex coacervation and slows molecular exchange within R1 condensates. (A) Schematic comparison of complex coacervation driven by charge–charge interactions and simple coacervation driven by aromatic and hydrophobic interactions. (B) Visual appearance of R1-FITC before and after addition of poly(U), and after centrifugation (200 μM R1, 5% FITC-R1, 200 μg/mL poly(U)). (C) Workflow for separating the condensed phase from the supernatant by centrifugation. (D) Quantification of poly(U) concentration and turbidity in the initial mixture, supernatant, and region above the pellet for control, R1, and R5 samples in D_2_O and H_2_O. (E) Turbidity profiles of R1/poly(U) mixtures in H_2_O and D_2_O as a function of poly(U) concentration (30-100 μM peptides, variable μg/mL poly(U)). (F) Representative FRAP recovery curves for R1 condensates formed in H_2_O and D_2_O. (G) Representative fluorescence images of R1 condensates before bleaching, immediately after bleaching, and after recovery. Scale bars: 10 μm. (H) Circular dichroism spectra of R1 in the absence and presence of poly(U). Unless otherwise noted, LLPS experiments were performed in 25 mM HEPES at matched pH/pD 7.4 and 23 °C. Data are presented as the mean ± SD from three independent experiments. Statistical significance was determined using two-way ANOVA followed by Šídák’s multiple-comparisons test; ^****^p < 0.0001.

### Phase-separation studies of aromatic and glycosylated peptide variants

To examine the contribution of aromatic amino acids and the effects of D_2_O, we compared the RRASL derivatives RRFSL (**R2**) and RRWSL (**R3**) (Figure 3A-B). We found that D_2_O promoted a significantly higher turbidity response relative to H_2_O, with a sharper onset and higher maximum. This finding suggests that aromaticity plays a key role in stabilizing condensates, likely through π-π or cation-π interactions. A similar trend was observed for additional RRASL constructs carrying sugar modifications, for which turbidity values were consistently higher in D_2_O (Figure 3C-G). When we assessed the effect of sugar modifications, **R4**-α-GalNAc behaved similarly to unmodified **R1**, whereas **R7**, which carries three α-GalNAc sugar moieties, exhibited a marked reduction in turbidity. These findings indicate that higher levels of glycosylation strongly suppress LLPS. Comparison of the α- and β-linked glycopeptide variants in H_2_O further revealed distinct phase-separation behavior. Notably, the β-glucose-modified peptide **R6** exhibited a significantly lower LLPS propensity than the β-galactose-modified peptide **R5** (Figures 3E), demonstrating that even subtle differences in sugar identity can modulate condensate formation. Finally, the remarkable difference between **R7** and **R8** further supports a crucial contribution of aromatic residues to phase separation, even when these residues are modified with sugars. We next examined the dynamics of **R5**-**R7** condensates by FRAP (Figure 3H-3J). All three glycopeptides showed substantial recovery in both H_2_O and D_2_O, with only modest, construct-dependent differences. Thus, despite altering their turbidity profiles, D_2_O did not uniformly slow molecular recovery, indicating that condensate formation and internal dynamics respond differently to solvent-isotope substitution.

**Figure 3.**
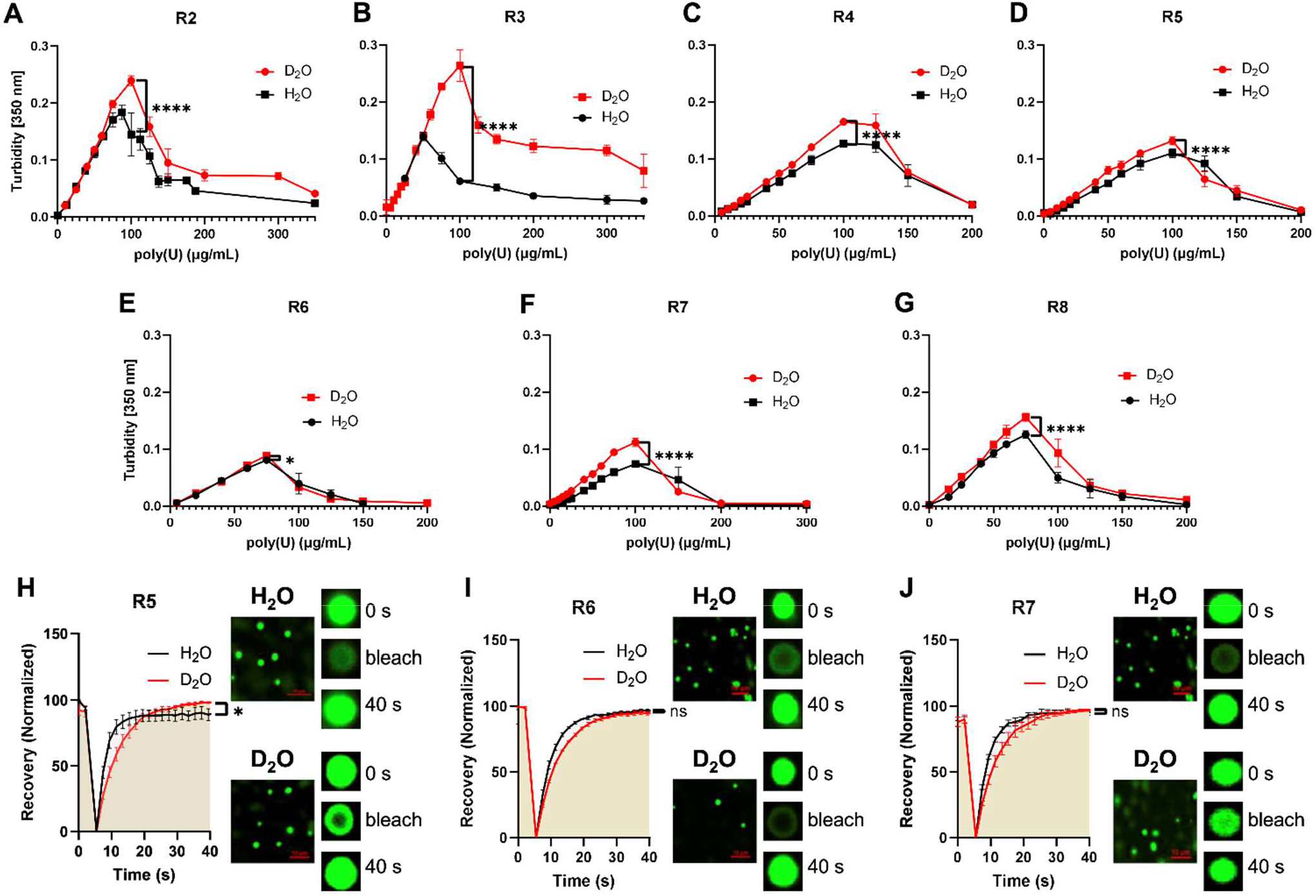
Aromatic residues and glycosylation tune solvent-isotope sensitivity in RRXSL peptide/poly(U) condensates. (A-G) Turbidity profiles of R2, R3, and glycopeptides R4-R8 mixed with variable concentrations of poly(U) in H_2_O and D_2_O. Peptide concentration was 100 μM in 25 mM HEPES at matched pH/pD 7.4 and 23 °C. (H-J) Representative FRAP recovery curves and fluorescence images for selected glycopeptide condensates formed in H_2_O and D_2_O. Scale bars: 10 μm. Data are presented as the mean ± SD or SEM for FREP from three independent experiments. Statistical significance between H_2_O and D_2_O conditions was determined using two-way ANOVA followed by Šídák’s multiple-comparisons test; ^*^p < 0.05, ^**^p < 0.01, ^****^p < 0.0001, and ns, not significant

### RNA-induced phase separation of tau constructs

Building on the results with peptide constructs, we next asked whether the solvent effects are applicable to intrinsically disordered proteins. Tau undergoes RNA-induced complex coacervation, and solvent isotope substitution exerts a substantial effect on both the effective RNA concentration required to reach maximal turbidity and the overall extent of phase separation (Figure 4). Turbidity measurements demonstrate that 2N4R tau, K18, and tau(291-391) all exhibit poly(U)-dependent phase separation characterized by reentrant behavior. For K18, the lysine-rich microtubule-binding domain shifts the RNA concentration required to reach maximal turbidity to ∼100 µg/mL in H_2_O, whereas in D_2_O the concentration required for maximal turbidity is reduced to ∼60 µg/mL, indicating enhanced interaction strength in heavy water (Figure 4A). A similar trend is observed for 2N4R tau, where turbidity modestly increases in D_2_O relative to H_2_O (Figure 4C). In contrast, for the shorter fragment tau(291-391), the concentration at which maximum turbidity occurs remains largely unchanged; however, the magnitude of turbidity increases sharply in D_2_O (Figure 4B).

**Figure 4.**
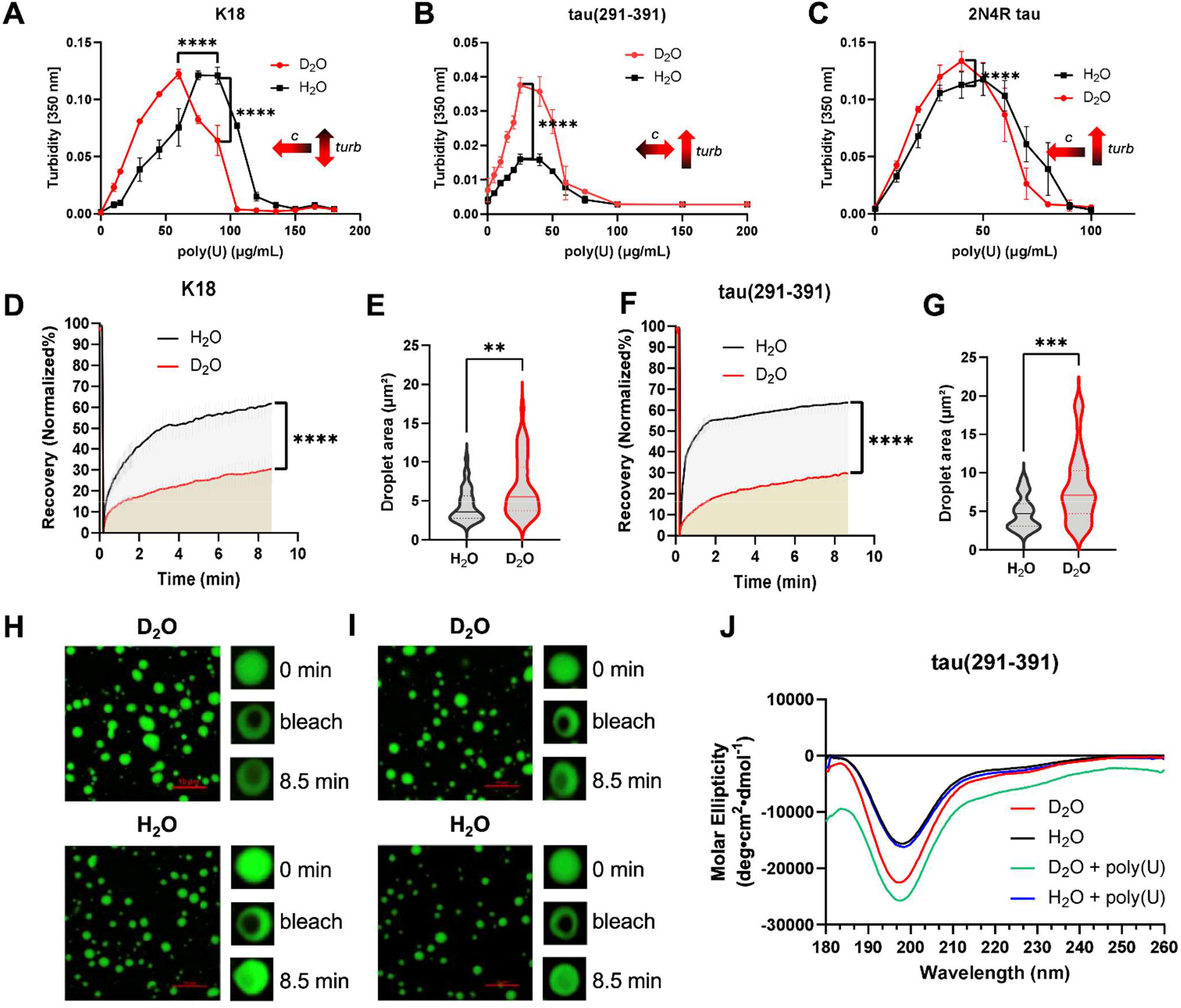
D_2_O shifts tau/RNA condensation and slows molecular exchange within tau condensates. (A–C) Turbidity profiles of K18, tau(291–391), and full-length 2N4R tau mixed with variable concentrations of poly(U) in H_2_O and D_2_O. LLPS assays were performed in 25 mM HEPES at matched pH/pD 7.4 and 23 °C. (D) FRAP recovery curves for K18 condensates formed in H_2_O and D_2_O. (E) Quantification of K18 condensate area under H_2_O and D_2_O conditions. (F) FRAP recovery curves for tau(291–391) condensates formed in H_2_O and D_2_O. (G) Quantification of tau(291–391) condensate area under H_2_O and D_2_O conditions. (H, I) Representative FRAP images of tau(291-391) and K18 condensates, respectively, before bleaching, immediately after bleaching, and after recovery. FRAP measurements were performed using solvent-specific protein and poly(U) concentrations corresponding to the maximum turbidity response and robust droplet formation in the respective H_2_O and D_2_O phase diagrams. Scale bars: 10 μm. (J) Circular dichroism spectra of tau(291–391) in the absence and presence of poly(U) in H_2_O and D_2_O. Data are presented as the mean ± SD from three independent experiments. Statistical significance for the turbidity and FRAP profiles was determined using two-way ANOVA followed by Šídák’s multiple-comparisons test, whereas droplet areas were compared using an unpaired two-tailed *t*-test; ^**^p < 0.01, ^***^p < 0.001, and ^****^p < 0.0001.

To further probe internal molecular dynamics, we performed FRAP experiments under both solvent conditions (Figure 4D-I). In H_2_O, droplets exhibited rapid fluorescence recovery, consistent with a highly dynamic, liquid-like internal environment. In contrast, droplets formed in D_2_O displayed significantly slower recovery kinetics and reduced molecular mobility. Quantitative analysis of recovery curves showed a statistically significant reduction in mobility in D_2_O. Confocal microscopy confirmed robust droplet formation in both solvents; however, droplet size distributions differed markedly (Figure 4H and 4I). Droplets formed in D_2_O were larger and more heterogeneous, whereas those in H_2_O were smaller and more uniformly distributed. Together, these results demonstrate that solvent isotope substitution alters the material properties of tau condensates by enhancing droplet stability and reducing internal molecular exchange. These findings suggest that strengthened hydrogen bonding in D_2_O modulates the physicochemical environment of tau, promoting a less dynamic condensate state. Notably, similar to the R1 peptide, LLPS did not significantly alter the secondary structure of tau within the condensates (Figure 4J).

To quantify the electrostatic interactions responsible for LLPS, we employed biolayer interferometry to directly measure the binding of K18 and tau(291-391) to RNA, using 5′-biotinylated U(40) immobilized on a streptavidin-coated biosensor probe. As shown in Figure 5, tau(291-391) in D_2_O exhibited a lower K_D_ value, 183 nM, compared with 712 nM in H_2_O but no significant differences for K18. These findings indicate that D_2_O enhances RNA-binding affinity, consistent with the enhanced LLPS observed in turbidity measurements (Figure 4). Together, the results suggest that the isotope effect of D_2_O arises from stronger protein-RNA interactions, which in turn promote more robust condensate formation.

**Figure 5.**
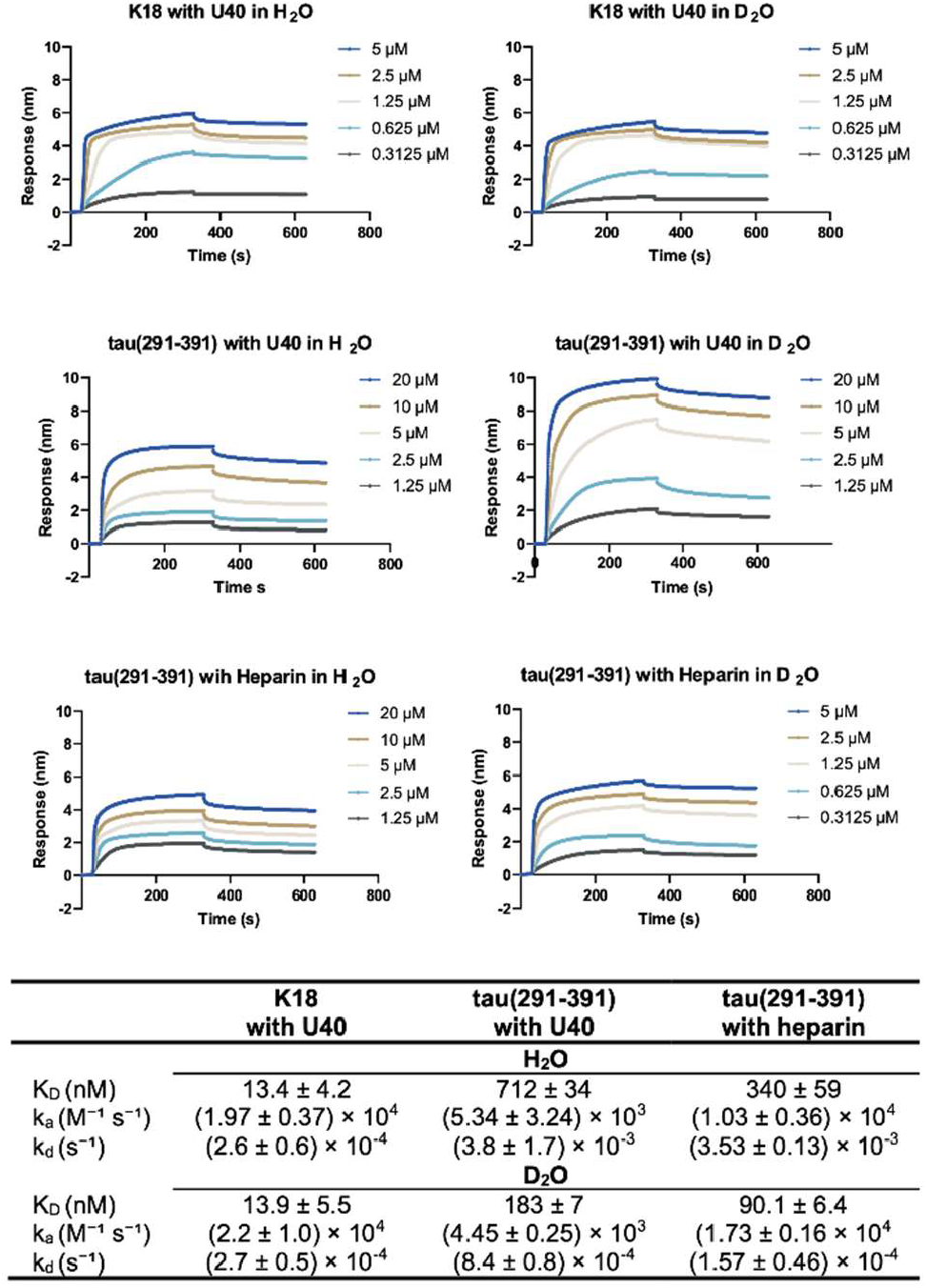
D_2_O selectively stabilizes labile tau-polyanion interactions. (A, B) Biolayer interferometry sensorgrams for K18 binding to immobilized U(40) RNA in H_2_O and D_2_O. (C, D) Sensorgrams for tau(291-391) binding to immobilized U(40) RNA in H_2_O and D_2_O. (E, F) Sensorgrams for tau(291-391) binding to immobilized heparin in H_2_O and D_2_O. U(40) RNA and heparin were immobilized on streptavidin biosensors, and tau proteins were applied as soluble analytes at the indicated concentrations. (G) Summary of apparent equilibrium and kinetic binding parameters, including K_D_, k_a_, and k_d_. Data are reported as mean ± SD from three independent measurements. Binding parameters were determined using a 1:1 binding model for U(40) and a 2:1 surface-avidity model for heparin.

### Salt-dependent phase-separation studie

To quantify how solvent isotope substitution alters the sensitivity of LLPS to ionic strength, we measured turbidity as a function of salt concentration (Figure 6). In H_2_O, increasing NaCl concentration led to a monotonic decrease in turbidity for all three systems, consistent with electrostatically driven phase separation. In contrast, condensates formed in D_2_O displayed substantially enhanced resistance to salt-induced dissolution in all systems examined. For (RRASL)_3_, turbidity in H_2_O decreased sharply with increasing NaCl and approached baseline by ∼30–40 mM NaCl (Figure 6A). In D_2_O, however, turbidity decayed more gradually and remained elevated across the entire salt range tested. Logistic fitting of the salt-response curves yielded an apparent half-maximal collapse concentration, IC_50,salt_, of ∼20 mM in H_2_O, whereas no collapse point was reached within the experimental window in D_2_O (IC_50,salt_ ≫ 40 mM). At the highest salt concentration tested, (RRASL)_3_ retained ∼35-40% of its initial turbidity in D_2_O compared with ∼8-10% in H_2_O.

**Figure 6.**
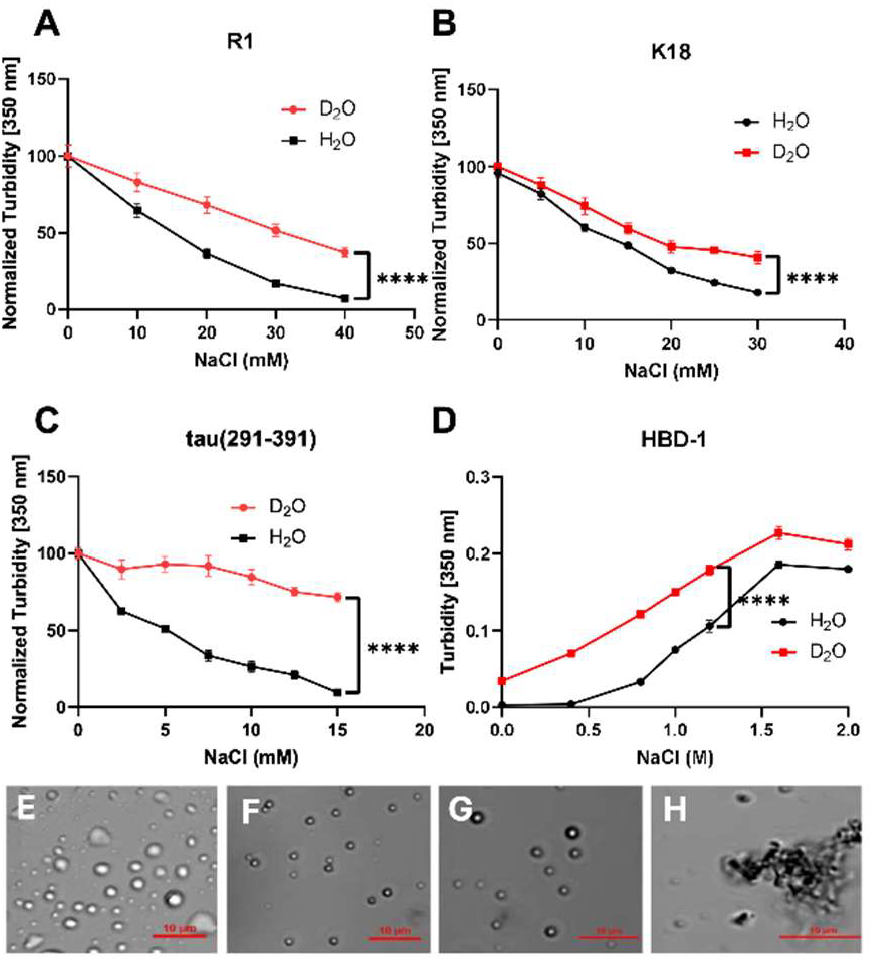
D_2_O stabilizes electrostatic condensates against salt-induced dissolution and alters salt-dependent R9/HBP-1 self-coacervation. (A-C) Salt-resistance assays for LLPS of (A) R1/poly(U), (B) K18/poly(U), and (C) tau(291-391)/poly(U) in H_2_O and D_2_O. Condensates were formed at the indicated peptide or protein and poly(U) concentrations and challenged with increasing NaCl in 25 mM HEPES at matched pH/pD 7.4 and 23 °C (100 μM R1, 100 μg·mL^−1^ poly(U); 60 μM tau(291-391), 30 μg·mL^-1^ poly(U); 60 μM K18, 60 μg·mL^-1^ poly(U)). Turbidity values are normalized to the no-salt condition. (D) Turbidity profile of the HBP-1-derived peptide R9 as a function of NaCl concentration in H_2_O and D_2_O. (E, F) Representative DIC images of R9 assemblies at 1 M NaCl in H_2_O and D_2_O, respectively. (G, H) Representative DIC images of R9 assemblies at 2 M NaCl in H_2_O and D_2_O, respectively. Scale bars: 10 μm. Data are presented as the mean ± SEM from three independent experiments. Statistical significance was determined using two-way ANOVA followed by Šídák’s multiple-comparisons test; ^****^p < 0.0001.

A similar but more pronounced trend was observed for tau-derived proteins (Figure 6B and 6C). For K18, LLPS in H_2_O was highly salt-sensitive, with a steep decline in turbidity at low mM NaCl concentrations. Substitution of H_2_O with D_2_O significantly shifted the salt-response curve upward, resulting in higher turbidity retention at all salt concentrations examined. The effect was most striking for tau(291-391) where turbidity collapsed almost completely by ∼10-15 mM NaCl in H_2_O, whereas in D_2_O, phase separation persisted across the entire salt range, with only a weak dependence on ionic strength. K18 showed a ∼2-3-fold increase in turbidity retention in D_2_O relative to H_2_O, while tau(291-391) exhibited the largest isotope effect, retaining >70% of its initial turbidity in D_2_O but approaching baseline in H_2_O. Analysis of the initial low-salt regime revealed that the slope of salt-induced turbidity decay was markedly reduced in D_2_O for all systems, approaching near zero for tau(291-391). This indicates that salt becomes a substantially weaker perturbation of intermolecular interactions in heavy water. Across all three systems, the magnitude of the solvent isotope effect correlated inversely with salt resistance in H_2_O. Condensates that were most sensitive to ionic screening in H_2_O, particularly tau(291-391), exhibited the strongest stabilization in D_2_O. These data indicate that heavy water does not merely increase baseline turbidity, but instead reshapes the phase boundary itself, shifting condensates from a marginal to a robust phase-separating regime under physiological ionic strengths (140-160 mM).^58^

We next characterized the morphology of HBP-1 under H_2_O and D_2_O conditions at different NaCl concentrations (Figure 6D). At 1 M NaCl, both solvents supported condensate formation, but at 2 M NaCl, the assemblies in D_2_O shifted completely from droplet-like structures to higher-order aggregates, whereas in H_2_O, condensates were still observed (Figure 6F and 6G). The D_2_O dependent shift from droplet-like condensates to higher-order aggregate suggests that solvent isotope substitution also modulates self-coacervation driven by hydrophobic and π-π stacking interactions, extending the isotope effect beyond electrostatically driven systems. Together results indicate that D_2_O has a pronounced impact on the aggregation behavior of proteins and peptides.

### Tau aggregation kinetics

Based on the observations on liquid-like assemblies, we postulated an alteration in amyloid-like aggregates in their fibrillation kinetics, fibril superstructure, and morphology (Figure 7). While the isotope effects have been reported for fibrillation of α-synuclein^59^ and insulin,^60^ the role of heavy water in tau aggregation is unknown. All three tau protein fragments exhibit the classical sigmoidal aggregation kinetics in both solvents.^61^ Fitting the ThT fluorescence kinetic curves to sigmoidal models allowed us to quantify the lag time, the half-time (t_½_), and an apparent rate constant for aggregation.^62, 63^ The K18 fragment in H_2_O displays a clear lag phase of roughly ∼10 h before ThT signal begins rising, followed by a rapid growth, and then a plateau/decline. In D_2_O, aggregation is faster, the lag phase is ∼1.5 h (about 3× shorter). Heavy water causes a ∼4-5× higher maximum ThT intensity (Figure S1A) for K18 (suggesting either more fibril formation or enhanced ThT binding) and an earlier plateau. The t_½_ for K18 drops from ∼17 h in H_2_O to ∼6.5 h in D_2_O (∼2.5× faster). Correspondingly, the apparent growth rate roughly doubles from ∼0.0059 min^™1^ in H_2_O vs ∼0.0093 min^™1^ in D_2_O.

**Figure 7.**
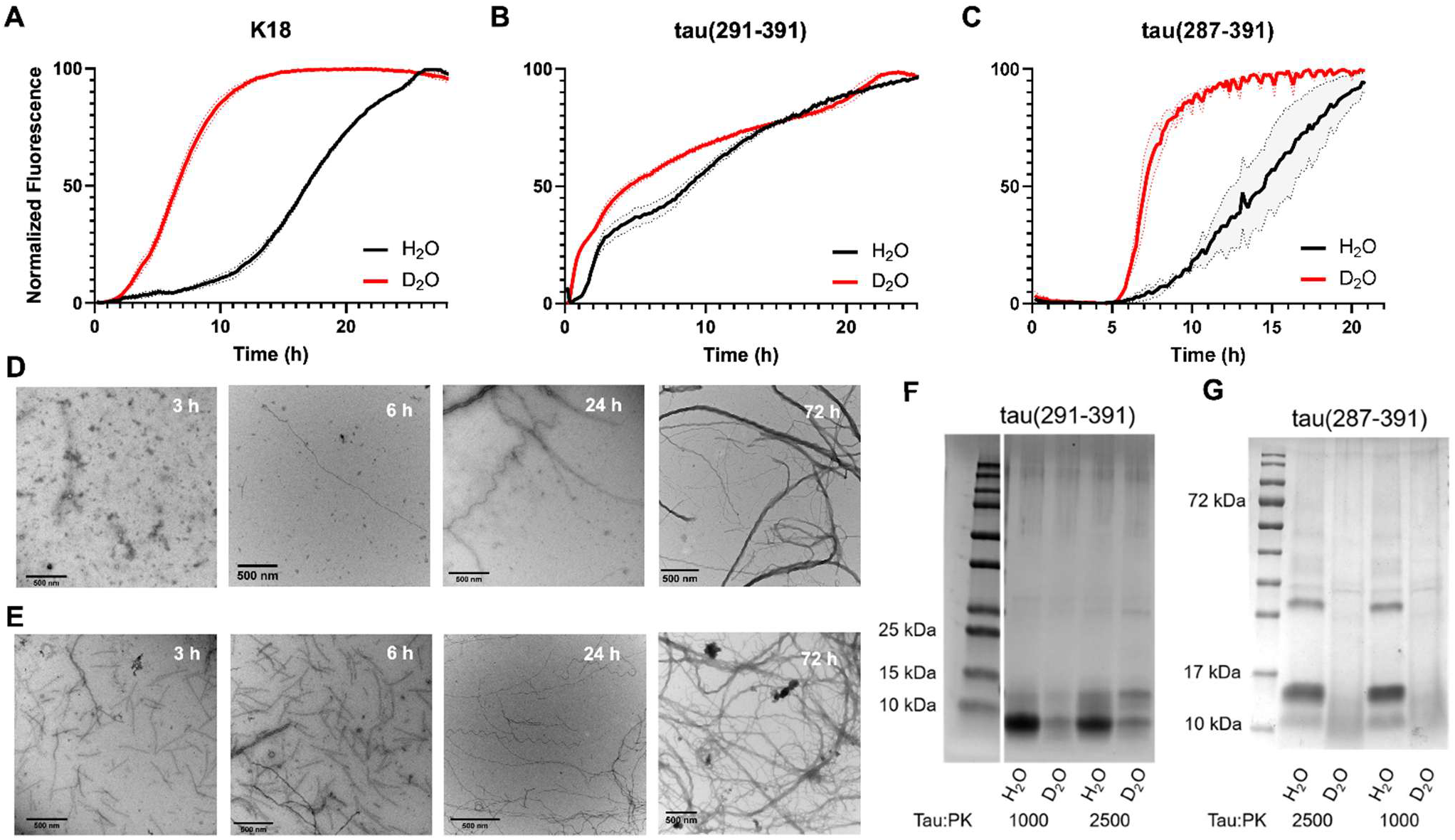
D_2_O accelerates tau amyloid formation and alters fibril organization. (A, B) Normalized fluorescence ThT aggregation kinetics of K18 and tau(291-391) in H_2_O and D_2_O under heparin-induced aggregation conditions (50 µM protein, 20 µM heparin, 10 mM DTT, 25 mM HEPES, 37 °C). (C) Cofactor-free normalized fluorescence ThT aggregation kinetics of tau(287-391) in H_2_O and D_2_O (6 mg/ml protein, 20 mM MgCl_2_, 10 mM DTT, 10 mM sodium phosphate, 200 RPM, 37 °C). (D, E) TEM time course of tau fibril formation in H_2_O and D_2_O at 3, 6, 24, and 72 h. (F, G) Proteinase K digestion analysis of tau(291-391) and tau(287-391) fibrils formed in H_2_O and D_2_O, followed by SDS-PAGE. Aggregation reactions were performed under matched pH/pD 7.4 conditions. Scale bars: 500 nm.

The tau(291-391) fragment showed the fastest aggregation (Figure 7B). Even in H_2_O, its ThT signal began rising almost immediately with only a very brief lag (∼1.5-2 h). The fluorescence increased steadily, reaching t_1/2_ by ∼14 h, and ultimately plateauing at ∼25 hours. In D_2_O, the kinetics were markedly accelerated, the lag phase was essentially negligible (≈1 h), the ThT intensity climbed much faster (t_1/2_ 5.3 h). Thus, for tau(291-391) as well, heavy water greatly speeds up aggregation, the half-time is about 2.6× shorter, and the maximum ThT signal ∼1.5× higher in D_2_O. In H_2_O, tau(287-391) showed an extended lag of ∼9 h, then a gradual increase in ThT fluorescence that reached its plateau only by ∼20-22 h. The sigmoidal curve here is less steep, suggesting a slower fibril growth phase. In D_2_O, we observe a faster aggregation, the lag phase shortened to ∼6 h, and the growth phase commenced earlier. The half-time t_½_ dropped from ∼14 h (H_2_O) to ∼7.3 h in D_2_O. The ∼4-fold increase in apparent tau(291-391)– heparin affinity in D_2_O, driven primarily by an ∼2.2-fold slower dissociation rate, increased the estimated lifetime of the bound complex from ∼5 to ∼11 min (Figure 5). This stabilization is consistent with the shorter aggregation lag phase and lower aggregation half-time observed in D_2_O, suggesting that long-lived tau–heparin complexes promote formation of nucleation-competent assemblies.

### Mechanical stability and proteolytic susceptibility

Based on the distinct aggregation kinetics and final ThT intensity, we hypothesized different final fibril amounts. To compare the extent of fibril formation in both solvents, tau proteoforms were aggregated under identical conditions followed by centrifugation, and the remaining soluble protein in the supernatant was quantified using a BCA assay. Across all tau proteoforms examined, the amount of soluble tau remaining after aggregation was consistently higher in H_2_O than in D_2_O, with the concentration ratios of 1.59:1 for tau(287-391), 1.77:1 for K18, and 1.82:1 for tau(291-391). Next, we employed transmission electron microscopy (TEM) to directly visualize the species that formed during the course of aggregation in both solvents to support our previous observations (Figure 7D and 7E). In H_2_O, only small oligomeric or amorphous assemblies were observed at early time points (3-6 h), with detectable fibrillar structures appearing after 24 h and maturing into long, unbranched fibrils by 72 h. Conversely, in D_2_O, fibril formation was evident as early as 3-6 h, with abundant, well-defined fibrils emerging by 24 h and extensive, dense fibrillar networks visible at 72 h.^64^

Additional characterization of the fibrils revealed solvent-dependent differences in stability and supramolecular organization. To evaluate the fibrils under mechanical stress, tau fibrils generated in both solvents were subjected to sonication followed by TEM analysis. Prior to sonication, fibrils formed in D_2_O appeared more bundled and interconnected relative to those formed in H_2_O. Following sonication, fibrils formed in H_2_O underwent extensive fragmentation into shorter and more dispersed species, whereas D_2_O-derived fibrils remained comparatively intact with fewer small fragments observed, indicating greater resistance to mechanical fragmentation.

To further probe structural differences, fibrils were treated with proteinase K followed by SDS-PAGE analysis (Figure 7F and 7G). Proteinase K is a non-specific enzyme that digests solvent-accessible regions and therefore serves as a probe for fibril structural compactness. Interestingly, fibrils formed from both tau(291-391) and tau(287-391) in H_2_O showed greater resistance to digestion compared with fibrils formed in D_2_O, as evidenced by the persistence of partially digested bands around 10 kDa. In contrast, D_2_O-derived fibrils exhibited reduced band intensity following digestion, suggesting increased proteolytic accessibility.

### Cellular seeding activity

Based on our proteinase K susceptibility and mechanical stability data that indicated alternative supramolecular packing for H_2_O and D_2_O raised fibrils, next we asked that whether this supramolecular features translate into alternative seeding competence in HEK293 biosensor cells via templated propagation recruiting tau monomers.^65, 66^ A HEK293T fluorescence resonance energy transfer (FRET) biosensor cell line that detects and amplifies seeding-competent tau aggregates provides a sensitive test for structural changes and overall seeding competency.^67^ In the absence of high-resolution structural data, this assay can serve as a proxy to evaluate structural differences. The biosensor cells were exposed to K18 and tau(291-391) fibrils formed under the above conditions, and the seeding efficiency was quantified by flow cytometry (FACS) based on the fraction of cells showing the GFP-YFP FRET signal (Figure 8). The Tukey post hoc analysis (family-wise α = 0.05) showed that tau(291-391) fibrils formed in H_2_O and D_2_O both produced significantly higher fluorescence intensities than the vehicle control (p < 0.0001 and p = 0.0001, respectively). Furthermore, tau(291-391) fibrils formed in H_2_O displayed significantly higher seeding activity than those formed in D_2_O (p = 0.0057). In the case of K18, fibrils formed in H_2_O and D_2_O also exhibited significantly greater seeding activity than the vehicle control (p = 0.0001 and p = 0.0044, respectively). Although K18 fibrils formed in D_2_O remained highly active seeds, their seeding activity was significantly reduced relative to K18 fibrils formed in H_2_O (p = 0.0388).

**Figure 8.**
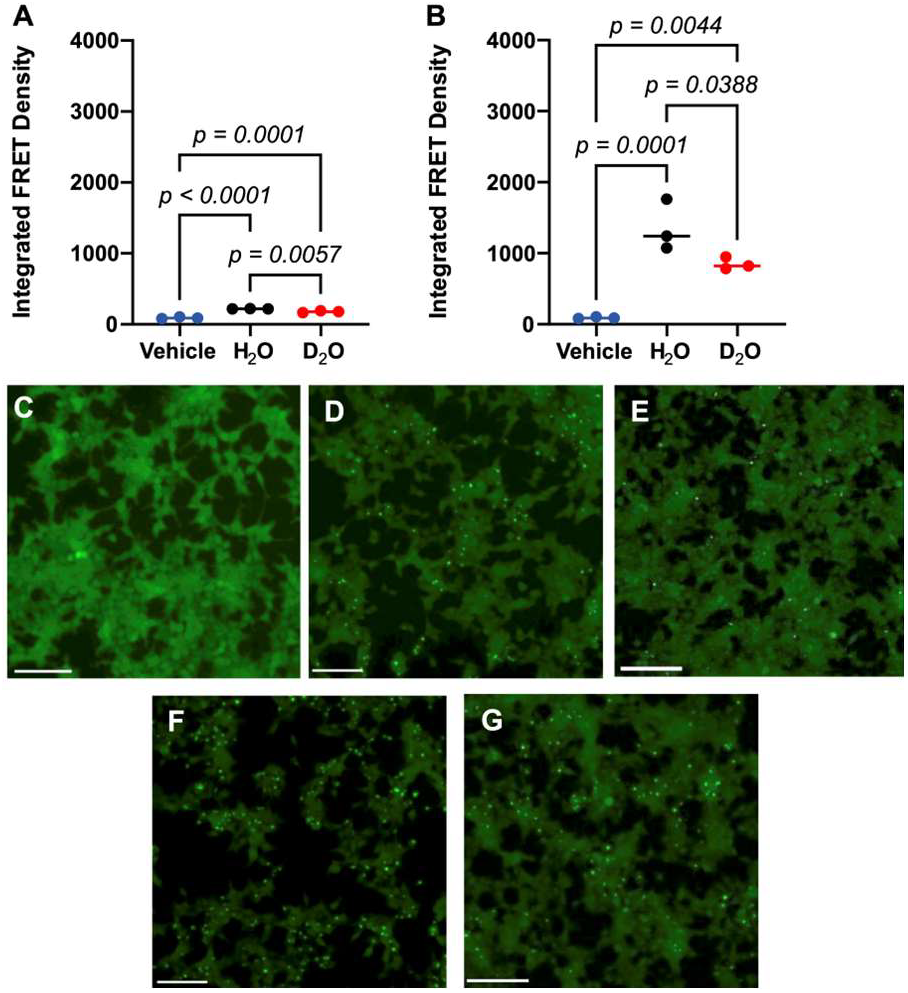
D_2_O-derived tau fibrils show altered seed competence in tau biosensor cells. (A-B) Quantification of integrated FRET density for tau(291-391) (A) and K18 (B) fibrils formed in H_2_O or D_2_O. Vehicle control in both panels are the same datasets reproduced for clarity. (C-G) Representative HEK293T biosensor images after treatment with vehicle (C), tau(291-391) fibrils formed in H_2_O or D_2_O (D-E), and K18 fibrils formed in H_2_O or D_2_O (F-G). Scale bars: 100 μm. Data are shown as individual values from n = 3 independent experiments. Statistical significance was determined using two-way ANOVA followed by Šídák’s multiple-comparisons test. Exact p values are indicated above the corresponding comparisons.

## DISCUSSION

Solvent isotope substitution is often treated as a technical adjustment rather than as a perturbation of biomolecular self-assembly. The results presented here support a solvent-isotope threshold model in which heavy water acts as a hydration-dependent amplifier of marginal biomolecular self-assembly. Rather than creating a new assembly pathway, D_2_O shifts pre-existing equilibria toward more associated and salt-resistant states, with construct-dependent effects on molecular dynamics. This effect was observed across chemically distinct systems, including His/Arg-rich peptides, glycopeptides, IDP coacervates, and amyloid assemblies. Importantly, the response was not uniform, and D_2_O had the largest effects on systems that were weakly associated, salt-sensitive, or close to an assembly boundary in H_2_O, whereas highly hydrated or already high-affinity systems showed smaller or qualitatively different responses. In peptide/RNA and tau/RNA condensates, D_2_O appears to lower the threshold for productive multivalent association, but in tau aggregation assays, this same principle manifests as shortened lag times and greater conversion into insoluble assemblies. However, D_2_O did not uniformly slow fluorescence recovery across all constructs. Notably, the glycopeptide condensates exhibited only modest, construct-dependent changes in FRAP recovery. This distinct behavior underscores that the solvent-isotope response is not governed by bulk viscosity alone but emerges from the interplay among viscosity, hydration, and intermolecular interactions, with peptide sequence and glycan composition determining their relative contributions.

The salt-resistance experiments provide the operational predictor of D_2_O responsiveness. Electrostatically driven complex coacervates were all destabilized by NaCl in H_2_O, but in D_2_O, these condensates retained substantially more turbidity over the same salt range. This relationship suggests that H_2_O salt sensitivity reports how close a condensate is to its dissolution boundary and can therefore predict the magnitude of the D_2_O response. Condensates that are already robust in H_2_O may still change morphology or dynamics in D_2_O, but marginal condensates should be strongly stabilized.

Biolayer interferometry supports this threshold model at the level of defined tau-polyanion interactions. K18 bound RNA with high apparent affinity in H_2_O, and this interaction changed little in D_2_O. In contrast, tau(291-391) showed a pronounced D_2_O-dependent increase in apparent affinity for RNA, and an even larger increase for heparin. These changes were driven primarily by slower dissociation rates in D_2_O, indicating stabilization of selected bound states rather than a uniform increase in all tau-polyanion interactions. This distinction may explain why K18 and tau(291-391) respond differently: the K18-U(40) interaction is already near a high-affinity limit under the assay conditions, whereas tau(291-391) interactions remain more labile and therefore more sensitive to solvent isotope substitution.

The same threshold principle extends from liquid-like condensates to amyloid formation. D_2_O accelerated aggregation of tau, indicating that solvent isotope substitution lowers the barrier for tau self-association under both heparin-induced and cofactor-free conditions. The tau(287-391) result is particularly informative because this construct aggregates without heparin, demonstrating that the D_2_O effect is not limited to enhanced tau-polyanion binding. However, faster aggregation did not translate into greater biological seed competence. K18 fibrils formed in D_2_O aggregated rapidly and remained seeding competent, but they produced a weaker biosensor response than K18 fibrils formed in H_2_O. These data support a seed competence model in which amyloid formation and functional templating activity are separable outputs. This model reconciles the aggregation, sonication, proteinase K, and biosensor results. D_2_O-derived fibrils appeared more bundled and were more resistant to mechanical fragmentation, suggesting increased supramolecular cohesion or network-level association. At the same time, D_2_O-derived fibrils were more sensitive to proteinase K digestion, indicating greater local protease accessibility. Therefore, reduced seeding by D_2_O-derived K18 fibrils should not be interpreted as evidence for a more compact or more protected fibril core, but, instead, D_2_O appears to alter fibril organization, producing assemblies that form readily but expose fewer productive templating units, fragment less efficiently, or present surfaces that are less compatible with intracellular tau recruitment.

In summary, D_2_O acts as a hydration-dependent amplifier of marginal biomolecular self-assembly. It stabilizes electrostatic condensates, slows molecular exchange, strengthens tau-polyanion interactions, and accelerates tau amyloid formation. The central conclusion is that solvent isotope substitution reshapes the energetic landscape of biomolecular self-assembly in a sequence-, modification-, and pathway-dependent manner. In tau, this produces a particularly important uncoupling as D_2_O accelerates amyloid formation but does not increase seed competence.

## ACKNOWLEDGMENT

This work was supported by the NIH (RF1AG079294, R01AG087295, R21GM138809). Mass spectrometry analyses were performed at the Proteomics and Mass Spectrometry Shared Research Core Facility at the University of Colorado Boulder (RRID: SCR_018992), which was funded by NIH grant S10OD038278. The imaging work was performed at the BioFrontiers Institute’s Advanced Light Microscopy Core (RRID: SCR_018302) and Flow Cytometry Shared Core (RRID:SCR_019309). We thank the Shared Instruments Pool (RRID: SCR_018986) at the University of Colorado Boulder for the use of the Applied Photophysics Chirascan Plus CD spectrometer which was funded by NIH S10RR028036. Electron microscopy was done in the EM Services Core Facility in the MCDB Department at the University of Colorado Boulder (RRID:SCR_001432). Figures were generated with the assistance of biorender.com.

