## Supplementary File for "Solvent-Isotope Effects in Biomolecular Phase Separation and Fibrillation of Disordered Proteins and Peptides"

### 1. General Information

All reactions were performed in oven-dried glassware. Reagents and solvents were obtained from commercial sources and used as received. All peptide coupling reactions were carried out in reactor vials. Peptide resins were swollen in *N,N*-dimethylformamide (DMF) before synthesis. Solid-phase peptide synthesis (SPPS) was carried out on a Biotage Initiator + Alstra™ microwave peptide synthesizer following the manufacturer's protocols unless otherwise noted. Preparative high-performance liquid chromatography (HPLC) was conducted using a Teledyne ISCO ACCQ Prep HP125 system. Liquid chromatography–mass spectrometry (LC–MS) analyses were performed on an Agilent 1260 Infinity II HPLC coupled to an Agilent MSD mass spectrometer or on an Advion Expression L instrument equipped with positive-mode electrospray ionization (ESI+). Mass spectra were acquired under standard operating conditions and deconvoluted using the UniDec graphical user interface (GUI-UniDec) software. Aqueous buffers were filtered through 0.22 μm syringe filters prior to use. UV absorbance at 280 nm and turbidity measurements were obtained using a NanoDrop 2000 spectrophotometer (Thermo Scientific). Microplate assays were conducted using a SpectraMax iD5 plate reader (Molecular Devices). Biolayer interferometry experiments were performed on a Sartorius Octet N1 system according to the manufacturer's protocols. Circular dichroism (CD) spectra were recorded on a Chirascan Plus CD spectrometer (Applied Photophysics). Negative-stain transmission electron microscopy (TEM) images were collected on an FEI Tecnai T12 Spirit microscope operated at 120 kV with a $LaB_6$ filament. Optical microscopy images were acquired using a Nikon Eclipse Ti laser-scanning confocal microscope equipped with a 60× oil-immersion objective. Image analysis was performed using ImageJ, and graphical representation and statistical analyses were carried out using GraphPad Prism.

All unmodified peptides, including **R1–R3** and **R9-HBP-1**, were purchased from GenScript and used as received.

### 2. Detailed Synthetic Procedures

#### 2.1. Peptide R4 (SPPS)

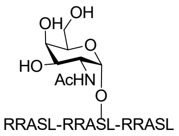

**R4**

**α-GalNAc-(RRASL)$_3$**

**Sequence:** RRASLRRA(α-GalNAc) SLRRASL-OH.
**Resin:** HMPB-MBHA resin from Sigma-Aldrich (100-200 mesh, 0.61 mmol/g, 410 mg, 0.25 mmol) was used. The resin was loaded by the symmetrical anhydride with Fmoc-L-Leu-OH, DIC, and 4- DMAP.
**Amino Acids:** Fmoc-L-Ser((Ac)$_3$-α-D-GalNAc)-OH, Fmoc-L-Ser($^t$Bu)-OH, Fmoc-L-Leu-OH, Fmoc-L-Arg(Pbf)-OH, Fmoc-L-Ala-OH.
**Coupling:** Amino acids were double coupled using 0.2 M amino acid in DMF (5 equiv.), 1 M DIC in DMF (10 equiv.), and 1 M oxyma pure in DMF (5 equiv.) at 75°C for 4 min. Fmoc-L-Ser((Ac)$_3$-α-D-GalNAc)-OH was coupled manually using 1.5 equiv. of the amino acid, 1.5 equiv. HATU, 1.5 equiv. Oxyma, and 5 equiv. DIPEA.
**Deprotection:** 20% piperidine + 0.1 M oxyma pure in DMF (6 mL) at 23 ºC (1 x 5 min, 1 x 15 min).
**Global Deprotection:** The crude resin-conjugated peptide was treated with TFA:$H_2O$:thioanisole:TIPSH (85:5:5:5, 20 mL) and stirred at 23 °C for 3 h. The reaction mixture was then cooled to 0 °C and treated with $Et_2O$ to precipitate the peptide. The precipitate was collected by centrifugation, and the resulting pellet was washed twice with $Et_2O$. The peptide was dissolved in 40% MeCN and lyophilized. Subsequently, the OAc protecting group was removed by treating the peptide with 4% $NH_2NH_2 \cdot H_2O$ at pH 9.5 and stirring at 23 °C for 3 h. The deprotected peptide was directly purified using a Luna C4 column (10 × 250 mm, 5 µm, 100 Å, 4.7 mL/min) with a 5–25% MeCN/$H_2O$/0.1% TFA gradient over 30 min. The collected fractions were lyophilized to afford **R4** (120 mg, 24% yield). Preparative HPLC trace of the crude deprotected peptide indicated that fractions 13–17 contained the desired peptide.

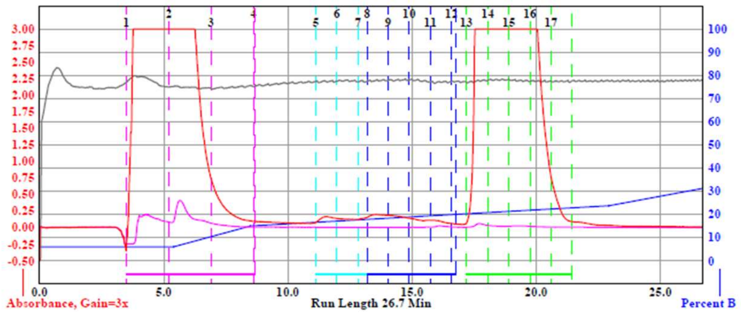

**Analytical LCMS:** Measured on an Agilent EC-C18 Poroshell column (4.6 x 100 mm, 4 µm, 100 Å, 40 °C, 1 mL/min) 5%-65% MeCN/$H_2O$/0.05% TFA over 30 min.
**Chemical Formula:** $C_{80}H_{150}N_{34}O_{24}$
**Molecular Weight:** 1971.29 g/mol

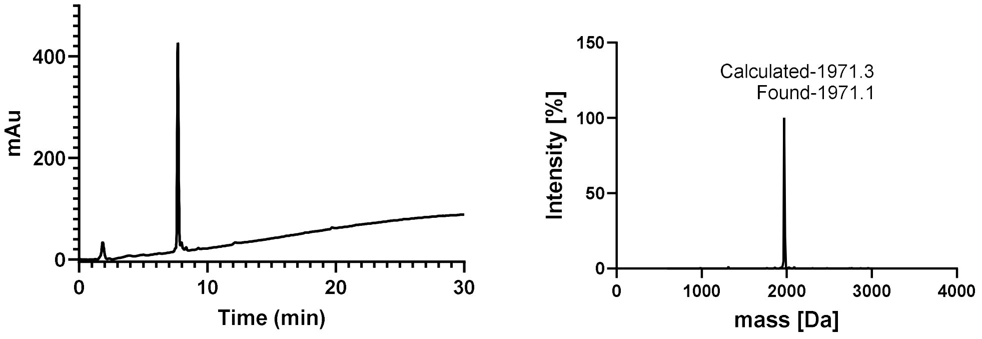

### 2.2. Peptide R5 (SPPS)

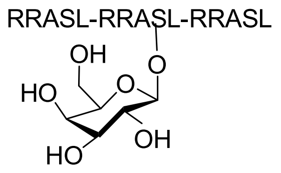

**R5**

**β-Gal-(RRASL)$_3$**

**Sequence:** RRASLRRA(β-Gal) SLRRASL-OH.
**Resin:** HMPB-MBHA resin from Sigma-Aldrich (100-200 mesh, 0.61 mmol/g, 410 mg, 0.25 mmol) was used. The resin was loaded by the symmetrical anhydride with Fmoc-L-Leu-OH, DIC, and 4- DMAP.
**Amino Acids:** Fmoc-L-Ser((Ac)$_4$-β-D-Gal)-OH, Fmoc-L-Ser($^t$Bu)-OH, Fmoc-L-Leu-OH, Fmoc-L-Arg(Pbf)-OH, Fmoc-L-Ala-OH.
**Coupling:** Amino acids were double coupled using 0.2 M amino acid in DMF (5 equiv.), 1 M DIC in DMF (10 equiv.), and 1 M oxyma pure in DMF (5 equiv.) at 75°C for 4 min. Fmoc-L-Ser((Ac)$_4$-β-D-Gal)-OH was coupled manually using 1.5 equiv. of the amino acid, 1.5 equiv. HATU, 1.5 equiv. Oxyma, and 5 equiv. DIPEA.
**Deprotection:** 20% piperidine + 0.1 M oxyma pure in DMF (6 mL) at 23 ºC (1 x 5 min, 1 x 15 min).
**Global Deprotection:** The crude resin-conjugated peptide was treated with TFA:$H_2O$:thioanisole:TIPSH (85:5:5:5, 20 mL) and stirred at 23 °C for 3 h. The reaction mixture was then cooled to 0 °C and treated with $Et_2O$ to precipitate the peptide. The precipitate was collected by centrifugation, and the resulting pellet was washed twice with $Et_2O$. The peptide was dissolved in 40% MeCN and lyophilized. Subsequently, the OAc protecting group was removed by treating the peptide with 4% $NH_2NH_2{\cdot}H_2O$ at pH 9.5 and stirring at 23 °C for 3 h. The deprotected peptide was directly purified using a Luna C4 column (10 × 250 mm, 5 µm, 100 Å, 4.7 mL/min) with a 5–26% MeCN/$H_2O$/0.1% TFA gradient over 35 min. The collected fractions were lyophilized to afford **R5** (134 mg, 27.8% yield). Preparative HPLC trace of the crude deprotected peptide indicated that fractions 21–33 contained the desired peptide.

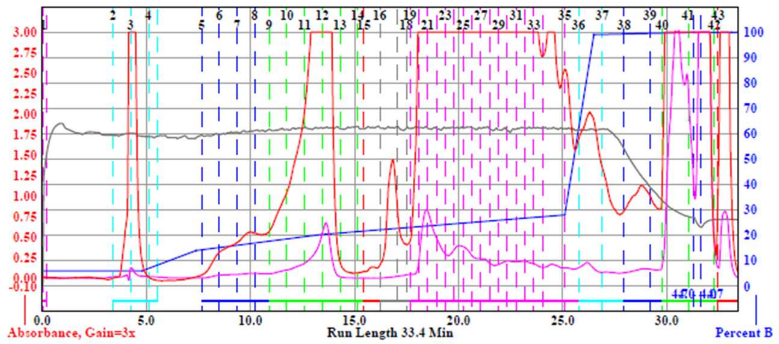

**Analytical LCMS:** Measured on an Agilent EC-C18 Poroshell column (4.6 x 100 mm, 4 µm, 100 Å, 40 °C, 1 mL/min) 5%-95% MeCN/$H_2O$/0.05% TFA over 30 min.
**Chemical Formula:** $C_{78}H_{147}N_{33}O_{24}$
**Molecular Weight:** 1931.24 g/mol

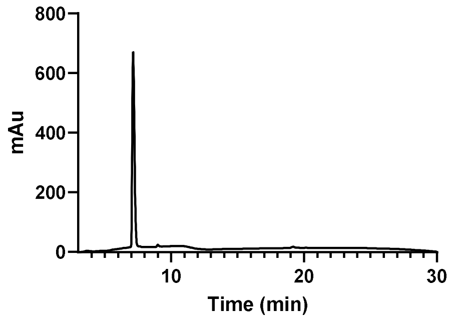

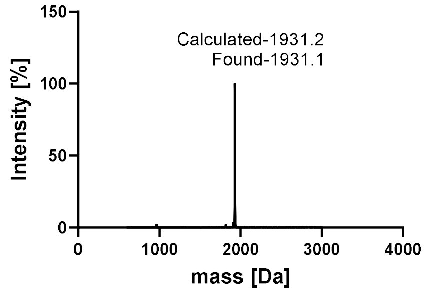

### 2.3. Peptide R6 (SPPS)

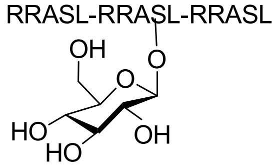

**R6**

**β-Glc-(RRASL)₃**

**Sequence:** RRASLRRA(β-Glc) SLRRASL-OH.
**Resin:** HMPB-MBHA resin from Sigma-Aldrich (100-200 mesh, 0.61 mmol/g, 410 mg, 0.25 mmol) was used. The resin was loaded by the symmetrical anhydride with Fmoc-L-Leu-OH, DIC, and 4- DMAP.
**Amino Acids:** Fmoc-L-Ser((Ac)$_4$-β-D-Glc)-OH, Fmoc-L-Ser($^t$Bu)-OH, Fmoc-L-Leu-OH, Fmoc-L-Arg(Pbf)-OH, Fmoc-L-Ala-OH.
**Coupling:** Amino acids were double coupled using 0.2 M amino acid in DMF (5 equiv.), 1 M DIC in DMF (10 equiv.), and 1 M oxyma pure in DMF (5 equiv.) at 75°C for 4 min. Fmoc-L-Ser((Ac)$_4$-β-D-Glc)-OH was coupled manually using 1.5 equiv. of the amino acid, 1.5 equiv. HATU, 1.5 equiv. Oxyma, and 5 equiv. DIPEA.
**Deprotection:** 20% piperidine + 0.1 M oxyma pure in DMF (6 mL) at 23 ºC (1 x 5 min, 1 x 15 min).
**Global Deprotection:** The crude resin-conjugated peptide was treated with TFA:$H_2O$:thioanisole:TIPSH (85:5:5:5, 20 mL) and stirred at 23 °C for 3 h. The reaction mixture was then cooled to 0 °C and treated with $Et_2O$ to precipitate the peptide. The precipitate was collected by centrifugation, and the resulting pellet was washed twice with $Et_2O$. The peptide was dissolved in 40% MeCN and lyophilized. Subsequently, the OAc protecting group was removed by treating the peptide with 4% $NH_2NH_2{\cdot}H_2O$ at pH 9.5 and stirring at 23 °C for 3 h. The deprotected peptide was directly purified using a Luna Phenyl-Hexyl (10 × 250 mm, 5 µm, 100 Å, 4.7 mL/min) with a 5–31% MeCN/$H_2O$/0.1% TFA gradient over 35 min. The collected fractions were lyophilized to afford **R6** (100 mg, 20.7% yield). Preparative HPLC trace of the crude deprotected peptide indicated that fractions 11–21 contained the desired peptide.

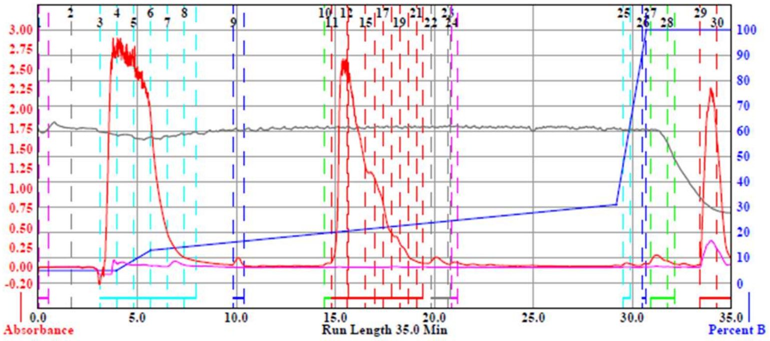

**Analytical LCMS:** Measured on an Agilent EC-C18 Poroshell column (4.6 x 100 mm, 4 µm, 100 Å, 40 °C, 1 mL/min) 5%-95% MeCN/$H_2O$/0.05% TFA over 30 min.
**Chemical Formula:** $C_{78}H_{147}N_{33}O_{24}$
**Molecular Weight:** 1931.24 g/mol

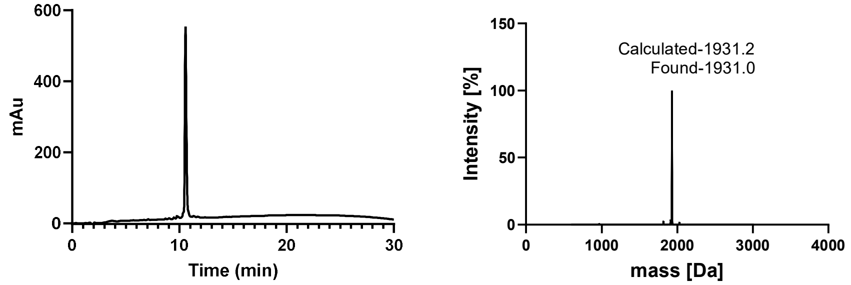

#### 2.4. Peptide R7 (SPPS)

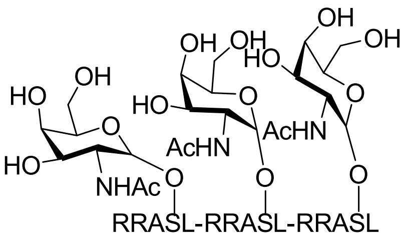

**R7**

**α-GalNAc$_3$-(RRASL)$_3$**

**Sequence:** RRASLRRA(α-GalNAc) SLRRASL-OH.
**Resin:** HMPB-MBHA resin from Sigma-Aldrich (100-200 mesh, 0.61 mmol/g, 163 mg, 0.1 mmol) was used. The resin was loaded by the symmetrical anhydride with Fmoc-L-Leu-OH, DIC, and 4- DMAP.
**Amino Acids:** Fmoc-L-Ser((Ac)$_3$-α-D-GalNAc)-OH, Fmoc-L-Ser($^t$Bu)-OH, Fmoc-L-Leu-OH, Fmoc-L-Arg(Pbf)-OH, Fmoc-L-Ala-OH.
**Coupling:** Amino acids were double coupled using 0.2 M amino acid in DMF (5 equiv.), 1 M DIC in DMF (10 equiv.), and 1 M oxyma pure in DMF (5 equiv.) at 75°C for 4 min. Fmoc-L-Ser((Ac)$_3$-α-D-GlcNAc)-was coupled manually using 1.5 equiv. of the amino acid, 1.5 equiv. HATU, 1.5 equiv. Oxyma, and 5 equiv. DIPEA.
**Deprotection:** 20% piperidine + 0.1 M oxyma pure in DMF (6 mL) at 23 ºC (1 x 5 min, 1 x 15 min).
**Global Deprotection:** The crude resin-conjugated peptide was treated with TFA:$H_2O$:thioanisole:TIPSH (85:5:5:5, 20 mL) and stirred at 23 °C for 3 h. The reaction mixture was then cooled to 0 °C and treated with $Et_2O$ to precipitate the peptide. The precipitate was collected by centrifugation, and the resulting pellet was washed twice with $Et_2O$. The peptide was dissolved in 40% MeCN and lyophilized. Subsequently, the OAc protecting group was removed by treating the peptide with 4% $NH_2NH_2 \cdot H_2O$ at pH 9.5 and stirring at 23 °C for 3 h. The deprotected peptide was directly purified using a Luna Phenyl-Hexyl (10 × 250 mm, 5 µm, 100 Å, 4.7 mL/min) with a 5–30% MeCN/$H_2O$/0.1% TFA gradient over 40 min. The collected fractions were lyophilized to afford **R7** (45 mg, 18.9% yield). Preparative HPLC trace of the crude deprotected peptide indicated that fractions 22–30 contained the desired peptide.

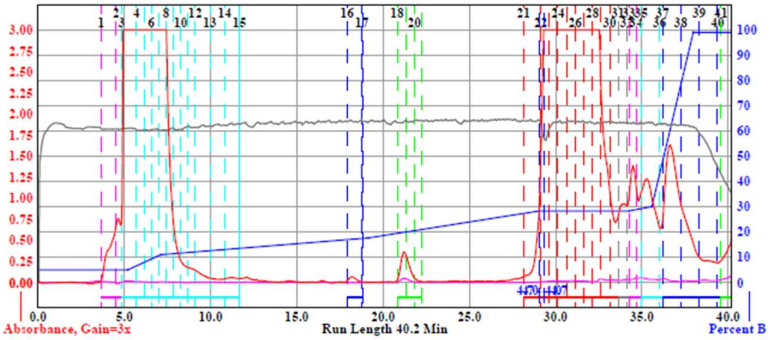

**Analytical LCMS:** Measured on an Agilent EC-C18 Poroshell column (4.6 x 100 mm, 4 µm, 100 Å, 40 °C, 1 mL/min) 5%-65% MeCN/$H_2O$/0.05% TFA over 30 min.
**Chemical Formula:** $C_{96}H_{176}N_{36}O_{34}$
**Molecular Weight:** 2378.68 g/mol

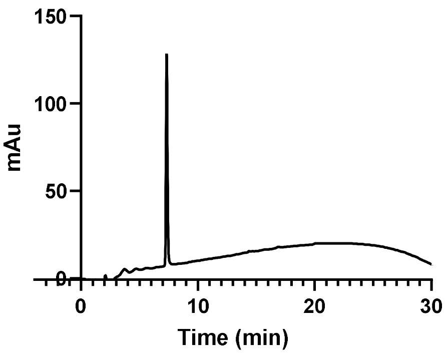

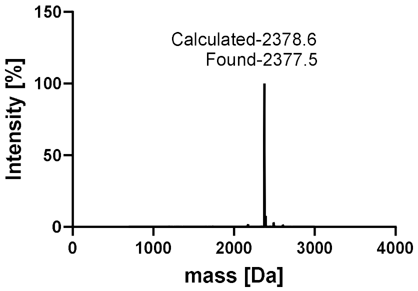

### 2.5. Peptide R8 (SPPS)

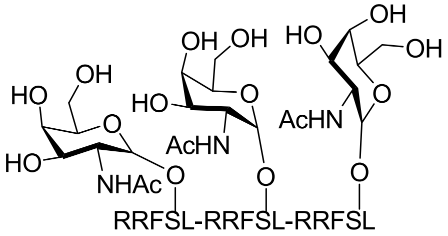

**R8**

**α-GalNAc$_3$-(RRFSL)$_3$**

**Sequence:** RRFSLRRF(α-GalNAc) SLRRFSL-OH.
**Resin:** HMPB-MBHA resin from Sigma-Aldrich (100-200 mesh, 0.61 mmol/g, 163 mg, 0.1 mmol) was used. The resin was loaded by the symmetrical anhydride with Fmoc-L-Leu-OH, DIC, and 4- DMAP.
**Amino Acids:** Fmoc-L-Ser((Ac)$_3$-α-D-GalNAc)-OH, Fmoc-L-Ser($^t$Bu)-OH, Fmoc-L-Leu-OH, Fmoc-L-Arg(Pbf)-OH, Fmoc-L-Phe-OH.
**Coupling:** Amino acids were double coupled using 0.2 M amino acid in DMF (5 equiv.), 1 M DIC in DMF (10 equiv.), and 1 M oxyma pure in DMF (5 equiv.) at 75°C for 4 min. Fmoc-L-Ser((Ac)$_3$-α-D-GlcNAc)-was coupled manually using 1.5 equiv. of the amino acid, 1.5 equiv. HATU, 1.5 equiv. Oxyma, and 5 equiv. DIPEA.
**Deprotection:** 20% piperidine + 0.1 M oxyma pure in DMF (6 mL) at 23 ºC (1 x 5 min, 1 x 15 min).
**Global Deprotection:** The crude resin-conjugated peptide was treated with TFA:$H_2O$:thioanisole:TIPSH (85:5:5:5, 20 mL) and stirred at 23 °C for 3 h. The reaction mixture was then cooled to 0 °C and treated with $Et_2O$ to precipitate the peptide. The precipitate was collected by centrifugation, and the resulting pellet was washed twice with $Et_2O$. The peptide was dissolved in 40% MeCN and lyophilized. Subsequently, the OAc protecting group was removed by treating the peptide with 4% $NH_2NH_2 \cdot H_2O$ at pH 9.5 and stirring at 23 °C for 3 h. The deprotected peptide was directly purified using a Luna Phenyl-Hexyl or Luna C4 column (10 × 250 mm, 5 µm, 100 Å, 4.7 mL/min) with a 5–35% MeCN/$H_2O$/0.1% TFA gradient over 35 min. The collected fractions were lyophilized to afford **R8** (53 mg, 20.3% yield). Preparative HPLC trace of the crude deprotected peptide indicated that fractions 18–22 contained the desired peptide.

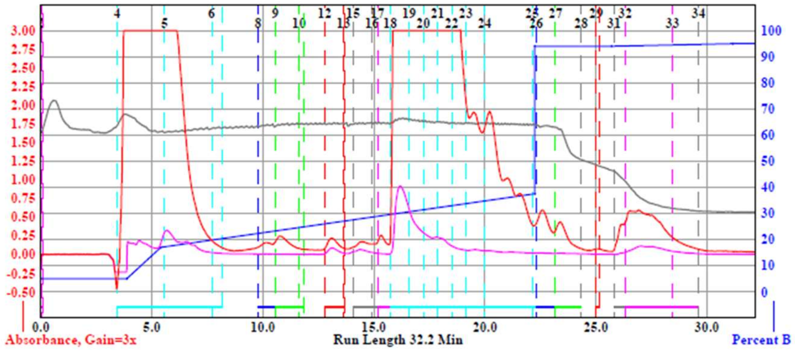

**Analytical LCMS:** Measured on an Agilent EC-C18 Poroshell column (4.6 x 100 mm, 4 µm, 100 Å, 40 °C, 1 mL/min) 5%-95% MeCN/$H_2O$/0.05% TFA over 30 min.
**Chemical Formula:** $C_{114}H_{188}N_{36}O_{34}$ **Molecular Weight:** 2606.97 g/mol.

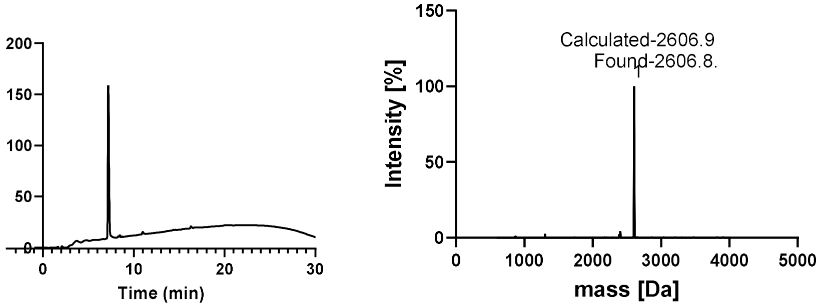

#### 2.6. Peptide tau(C291-E391) (SPPS)

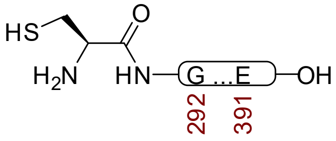

**tau(291-391)**

The synthesis of residues tau(291–391) was carried out following the protocol previously published.[1] In brief, the protein was synthesized using solid-phase peptide synthesis on a microwave-assisted synthesizer with Fmoc chemistry, followed by native chemical ligation. Amino acids were double coupled under the conditions described in the original protocol. After assembly, the crude resin-bound peptide was subjected to global deprotection using TFA-based cleavage, followed by precipitation with diethyl ether. The resulting peptide was purified by preparative HPLC, and fractions containing the desired peptide were collected and lyophilized.

The protein was further purified using sample displacement mode chromatography[2] on a Zorbax C18 column (4.6 × 250 mm, 5 µm, 300 Å) at 40 °C with a flow rate of 4.7 mL/min. A 5–65% MeCN/$H_2O$/0.05% TFA gradient was applied over 35 min to obtain the final pure peptide.

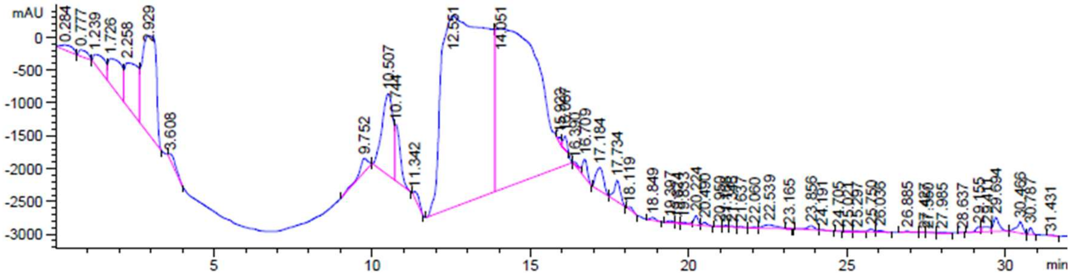

**Chemical Formula:** $C_{465}H_{764}N_{142}O_{147}S_2$
**Molecular Weight:** 10770.19 g/mol

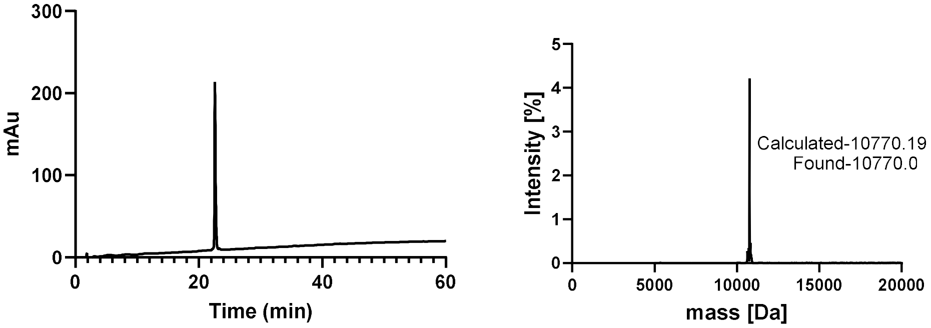

#### 2.7. Peptide tau(N244-E372) (SPPS)

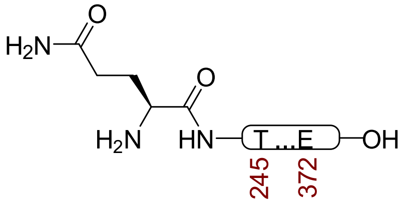

**K18**

The synthesis of residues K18 was carried out following the protocol previously described.[1] In brief, the protein was synthesized using solid-phase peptide synthesis on a microwave-assisted synthesizer with standard Fmoc chemistry, followed by native chemical ligation. Amino acids were double coupled under the conditions described in the original protocol. After assembly, the crude resin-bound peptide was subjected to global deprotection using TFA-based cleavage, followed by precipitation with diethyl ether. The resulting peptide was purified by preparative HPLC, and fractions containing the desired peptide were collected and lyophilized.

The material was further purified using sample displacement mode chromatography[2] on a Zorbax C18 column (4.6 × 250 mm, 5 µm, 300 Å) at 40 °C with a flow rate of 4.7 mL/min. A 5–65% MeCN/$H_2O$/0.05% TFA gradient was applied over 35 min to obtain the final pure peptide.

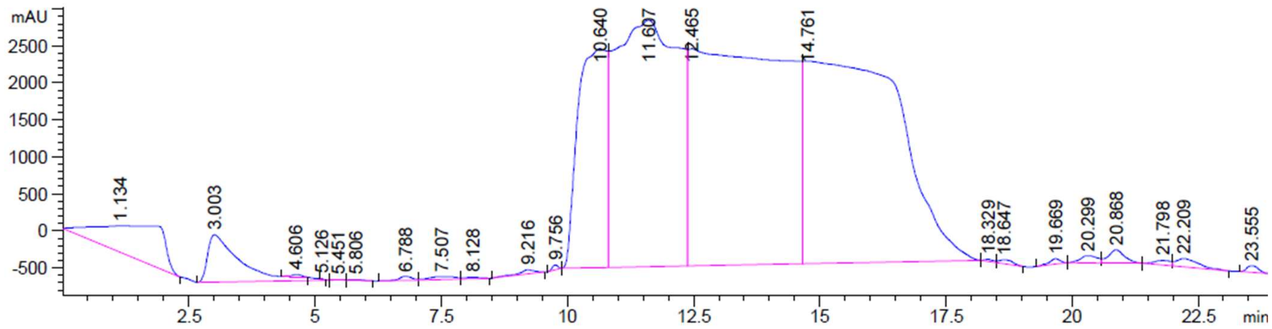

**Chemical Formula:** $C_{598}H_{1000}N_{178}O_{184}S_{3}$
**Molecular Weight:** 13723.1 g/mol

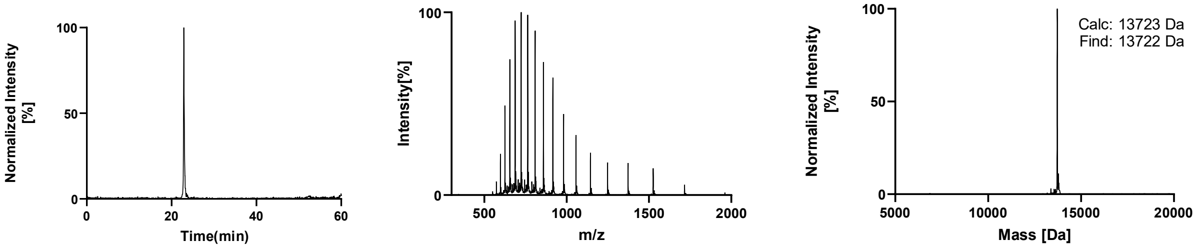

### 2.8. Recombinant Synthesis of tau(287-391)

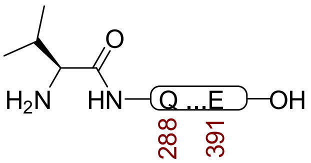

**tau(287-391)**

The tau(287-391) encoding sequence was cloned into the pET28b(+) vector by Genescript, and the plasmid was transformed into *E. coli* BL21 (DE3) cells. Overnight primary culture was set up from freshly transformed *E. coli* in lysogeny broth (LB) supplemented with kanamycin (50 µg/mL). 6 L of secondary culture was set up in LB with kanamycin (50 µg/mL) at 37 °C and 220 rpm until the culture reached an OD600 of 0.4-0.6. Then, the cultures were induced with 1 mM IPTG at 37 °C for 3 hours. Cells were harvested by centrifugation (4000× *g* for 25 min at 4 °C) and resuspended in washing buffer (WB: 50 mM MES, pH 6.0; 10 mM EDTA; 10 mM DTT, supplemented with Pierce protease cocktail inhibitors). Cell lysis was carried out using sonication (at 55 % amplitude using a Qsonica Sonic Processor for 12 min, 30 s on/30 s off) after incubating the cell suspension with 1.5 mg/ml lysozyme for 2 hours at 4 °C. Lysed cells were centrifuged at 12,000× *g* for 45 min at 4 °C, and the supernatant containing proteins was loaded onto two tandem CaptoS 5 ml columns (Cytiva) for cation exchange chromatography. The columns were washed with 10 column volumes of WB and eluted using a gradient of WB containing 0–1 M NaCl. Fractions of 1.9 ml were collected and analyzed by SDS-PAGE, and the protein-containing fractions were pooled and concentrated up to 2 ml using a 5 kDa cut-off Vivaspin protein concentrator unit (Sartorius). The concentrated protein was loaded into a HiLoad 16/600 Superdex 75 pg size exclusion column (Cytiva), and eluted in 10 mM sodium phosphate, pH 7.2–7.4, 10 mM DTT. Size exclusion fractions were analysed by SDS-PAGE, protein-containing fractions were pooled and concentrated to ~10 mg/ml using protein concentrators with a cut-off filter of 5 kDa, and flash frozen or lyophilized for storage. The protein was further purified using sample displacement mode chromatography[2] on a Zorbax C18 column (4.6 × 250 mm, 5 µm, 300 Å) at 40 °C with a flow rate of 4.7 mL/min. A 5–65%. MeCN/$H_2O$/0.05% TFA gradient was applied over 60 min to obtain the final pure peptide tau(287-391).

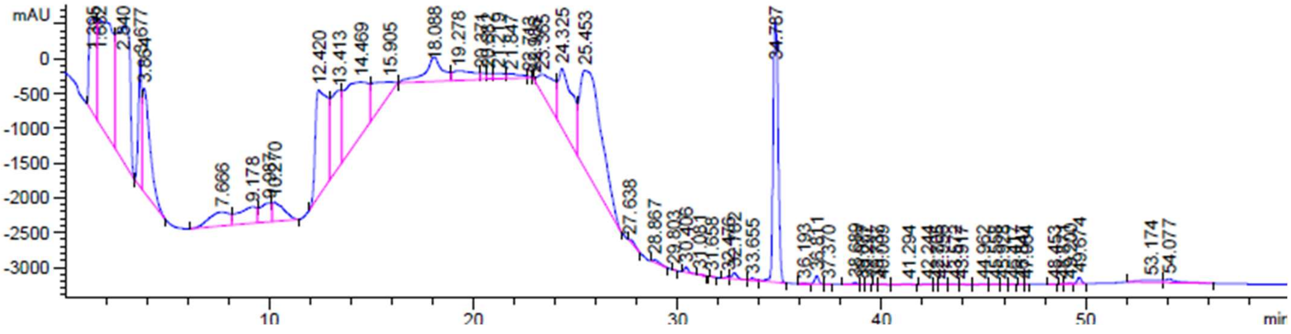

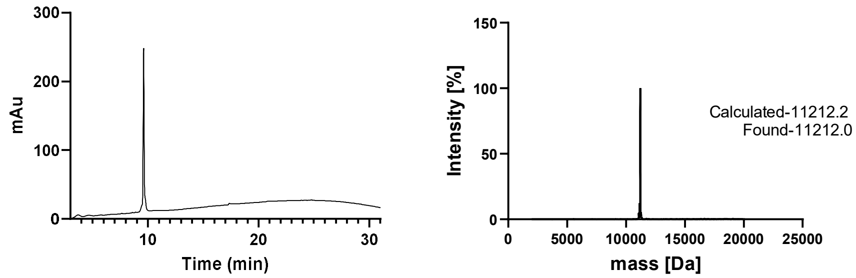

**Chemical Formula:** $C_{486}H_{801}N_{149}O_{151}S_2$
**Molecular Weight:** 11212.2 g/mol

### 2.9. Recombinant Synthesis of Full-Length 2N4R tau

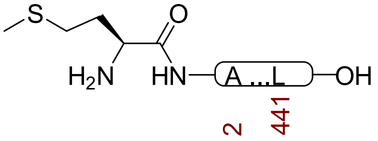

**2N4R tau**

Per published protocol,[3] N-terminal hexahistidine-SUMO tag with 2N4R tau encoding sequence was cloned into the vector pET30a(+) by Genescript, and the plasmid was transformed into *E. coli* BL21 (DE3) cells. An overnight primary culture was set up for His-SUMO-2N4R tau from freshly transformed *E. coli* in lysogeny broth (LB) supplemented with kanamycin (50 µg/mL). 4 L of secondary culture was set up in LB with kanamycin (50 µg/mL) at 37 °C and 220 rpm until the culture reached an OD600 of 0.4-0.6. Then, the cultures were induced with 1 mM IPTG at 37 °C for 3 hours. Cells were harvested by centrifugation (4000× *g* for 25 min at 4 °C) and resuspended in lysis buffer (25 mM HEPES, pH 7.4; 300 mM NaCl; 20 mM imidazole) supplemented with Pierce protease cocktail inhibitors. Cell lysis was carried out using sonication (at 55 % amplitude using a Qsonica Sonic Processor for 12 min, 30 s on/30 s off) after incubating the cell suspension with 1.5 mg/ml lysozyme for 2 hours at 4 °C. Lysed cells were centrifuged at 12,000× *g* for 45 min at 4 °C, and the supernatant containing proteins was mixed with lysis buffer-incubated Ni-NTA resin for 2 hours. Ni-IMAC was performed with a gradient of imidazole washes, and the protein was finally eluted using 280 mM imidazole supplemented in the lysis buffer. His-SUMO affinity tag was cleaved using the recombinant histidine-tagged SUMO protease Ulp1 (His-Ulp1). The cleavage reaction was purified by preparative HPLC using a Teledyne HP-150 system. The sample was loaded onto a Luna C4 column (10 × 250 mm, 5 µm, 100 Å, 4.7 mL/min) with a 5–35% MeCN/$H_2O$/0.1% TFA gradient over 35 min. Fractions were analyzed by LC/MSD on an Agilent EC-C18 Poroshell column (4.6 × 100 mm, 4 µm, 100 Å) at 40 °C and a flow rate of 1 mL/min. Fractions containing the desired product were combined and lyophilized at −80 °C to yield 15 mg of 2N4R tau as a white solid.

.

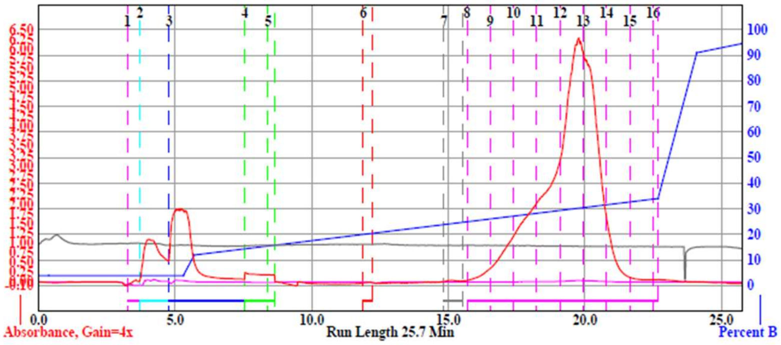

**Chemical Formula:** $C_{1959}H_{3197}N_{581}O_{669}S_8$
**Molecular Weight:** 45850.1 g/mol

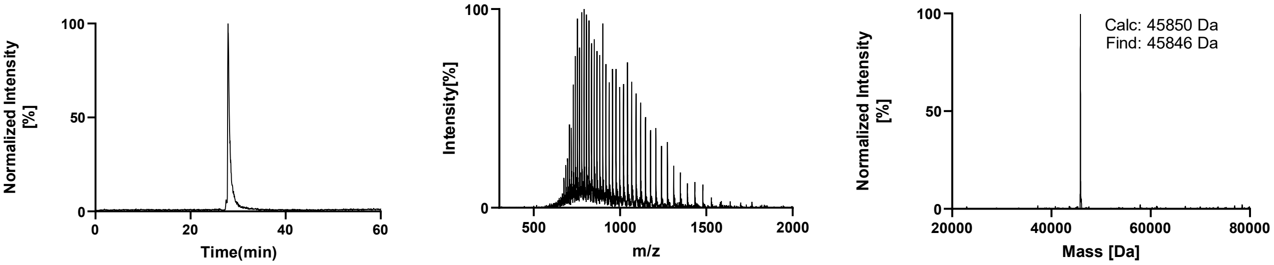

### 3. Protein Concentration Determination

Lyophilized Tau proteins were reconstituted in 25 mM HEPES buffer (pH/pD 7.4), and protein concentrations were determined by ultraviolet absorbance at 280 nm using a NanoDrop 1000 spectrophotometer. To prepare samples in both $H_2O$ and $D_2O$, a single peptide solution was divided into two aliquots, lyophilized, and one aliquot was redissolved in $D_2O$ and lyophilized again to allow proton–deuteron exchange. The final concentrations were then measured by NanoDrop after dissolved in 25 mM HEPES (pH/pD 7.4) either in $H_2O$ or $D_2O$.

In contrast, the concentrations of short synthetic peptides (R1–R9), which lack sufficient aromatic residues for reliable UV-based quantification, were determined gravimetrically. Lyophilized peptides were weighed using a balance, and concentrations were calculated based on their molecular weights. Each known weight of peptide was dissolved in a measured volume of $H_2O$, and the solution was divided into equal aliquots, both of which were lyophilized. To prepare $D_2O$ samples, one of the lyophilized aliquots was redissolved in $D_2O$ and lyophilized again to allow proton–deuteron exchange.

### 4. Deuterated Proteins and RNA Stock Preparations

For $D_2O$-based samples, all proteins, buffers, DTT, heparins, and poly(U) etc. were initially dissolved in $D_2O$. The solutions were frozen and lyophilized to remove residual $H_2O$. This dissolution and lyophilization cycle were repeated twice to ensure maximal exchange of labile protons and achieve efficient deuteration of all components. The final lyophilized materials were used for further experiments. Freshly deuterated samples were prepared for each experiment. For $D_2O$ samples, the pD 7.4 was maintained by adding NaOD or DCl solutions.

For $H_2O$-based samples, all proteins, buffers, DTT, heparins, and poly(U) were dissolved in $H_2O$ and subjected to the same freeze lyophilization procedure as used for the $D_2O$ samples to maintain identical sample processing conditions.

### 5. Co-Factor Free Aggregation

Lyophilized powder of tau(287-391) was dissolved in 10 mM Sodium Phosphate, pH/pD 7.4, 10 mM DTT, and the concentration was checked by absorption at 280. The final monomer concentration is 6 mg/mL, and the aggregation buffer consists of 10 mM Sodium Phosphate, pH/pD 7.4, 10 mM DTT, and 20 mM $MgCl_2$. The addition of compounds in reactions is preceded by first adding protein dissolved in 10 mM Sodium phosphate, pH/pD 7.4, and 10 mM DTT. Then, DTT and Sodium Phosphate were supplemented in the reactions to make a final concentration of 10 mM each, in addition to the Sodium phosphate, and DTT contributed from the protein stock. 20 mM $MgCl_2$ was added slowly to avoid precipitation. 20 µM ThT was added to the reactions to monitor aggregation kinetics using fluorescence at 480 nm. Aggregation reactions were set up in a VANTAstar Flexible Multi-mode Microplate Reader (BMG Labtech) using Corning 384 Well Black Polystyrene Microplate with orbital shaking at 200 rpm at 37°C with 6 nanobeads/well (800 µM Silica beads, OPS diagnostic). For the aggregation of deuterated tau(287-391), protein and buffer components were deuterated before setting up aggregation reactions, and the same process was followed.

### 6. Heparin-Induced Aggregation

Aggregation reactions (50 µL total volume per condition) containing tau(291–391) or K18 were mixed in 0.65 mL microcentrifuge tubes using either $D_2O$ or $H_2O$-based buffers at the final concentrations specified below. Buffer components and salts were combined first, followed by addition of the protein, with heparin introduced as the final component in all cases. Aliquots of the reaction mixture (15 µL) were dispensed into three separate wells of a black, non-binding 384-well microplate (Greiner Bio-One; polystyrene; flat-bottom; small volume; HiBase; REF 784900). The plate was sealed with a polyester adhesive film (VWR, Cat. No. 89134-430) and maintained at 37 °C in the microplate reader for fluorescence measurements.

Fluorescence signals from the three technical replicates were averaged for each condition. All experiments were performed in triplicate, and the normalized data are reported as mean ± standard error of the mean (SEM).

**Stock solutions were prepared either in $D_2O$ or $H_2O$ as follows:**
25 mM HEPES buffer (pH/pD 7.4).
100 mM DTT in 25 mM HEPES (pH/pD 7.4).
1 M NaCl in 25 mM HEPES (pH/pD 7.4).
10 mM heparin sodium salt from porcine intestinal mucosa (Sigma H4784; average molecular weight ~18 kDa; 1 mg/mL ≈ 55 µM) in 25 mM HEPES (pH/pD 7.4).
165 µM thioflavin T in 25 mM HEPES (pH/pD 7.4).
200–250 µM protein in 25 mM HEPES (pH/pD 7.4).

**Final assay conditions for both $D_2O$- and $H_2O$-based samples were:**

50 µM protein, 20 µM heparin, 10 µM thioflavin T, 10 mM DTT, 100 mM NaCl, and 25 mM HEPES (pH/pD 7.4), incubated at 37 °C.

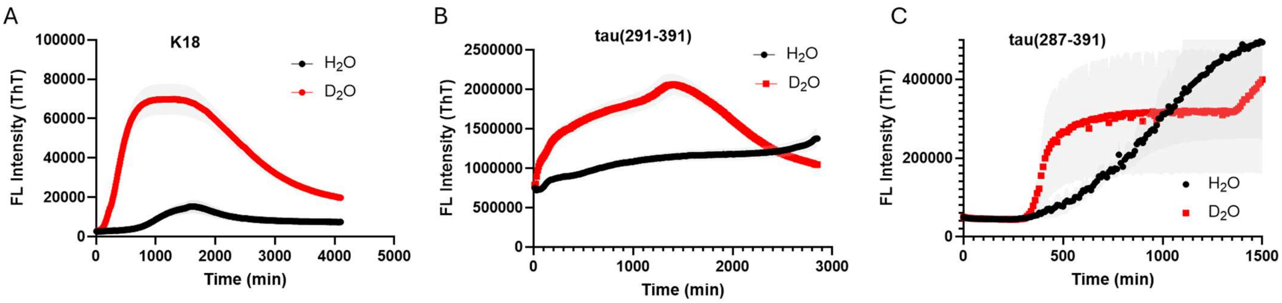

**Figure S1**. ThT aggregation kinetics of K18 and tau(291-391) in $H_2O$ and $D_2O$ under heparin-induced aggregation conditions (50 µM protein, 20 µM heparin, 10 mM DTT, 25 mM HEPES, 37 ºC). (C) Cofactor-free ThT aggregation kinetics of tau(287-391) in $H_2O$ and $D_2O$ (6 mg/ml protein, 20 mM $MgCl_2$, 10 mM DTT, 10 mM sodium phosphate, 200 RPM, 37 ºC).

### 7. Negative Staining TEM Procedures

Saturated aggregation reactions were diluted 10 times in water after ~ 4 days of commencement. 5 µL of diluted fibrils were applied to glow-discharged grids (Formvar/Carbon 200 Mesh, Copper) and incubated for 1 min, and extra samples were side-blotted. The grids were washed with 100 µL of filtered water. Then, 3.5 µL of 2% Uranyl acetate was applied to the grids, and the excess stain was completely side blotted after 1 min of incubation. The grids were air-dried for 5 min, and imaging was performed in the FEI Tecnai T12 Spirit Transmission Electron Microscope.

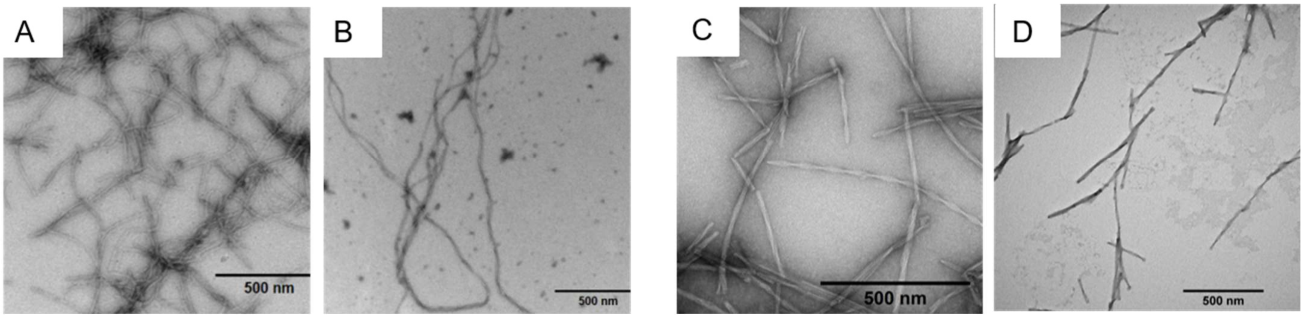

**Figure S2.** TEM images of fibrils formed after 72 h for K18 in $H_2O$ and $D_2O$ under heparin-induced aggregation conditions (50 µM protein, 20 µM heparin, 10 mM DTT, 25 mM HEPES (pH/pD 7.4), 37 °C. (A) K18 fibrils in $D_2O$ condition after 72 h (B) K18 fibrils in $H_2O$ condition after 72 h. Cofactor-free fibrillization conditions: 6 mg $mL^{-1}$ protein, 20 mM $MgCl_2$, 10 mM DTT, 10 mM sodium phosphate (pH/pD 7.4), 200 rpm, 37 °C. (C) tau(287–391) fibrils in $D_2O$ after condition 24 h. (D) Tau(287–391) fibrils in $H_2O$ condition after 24 h.

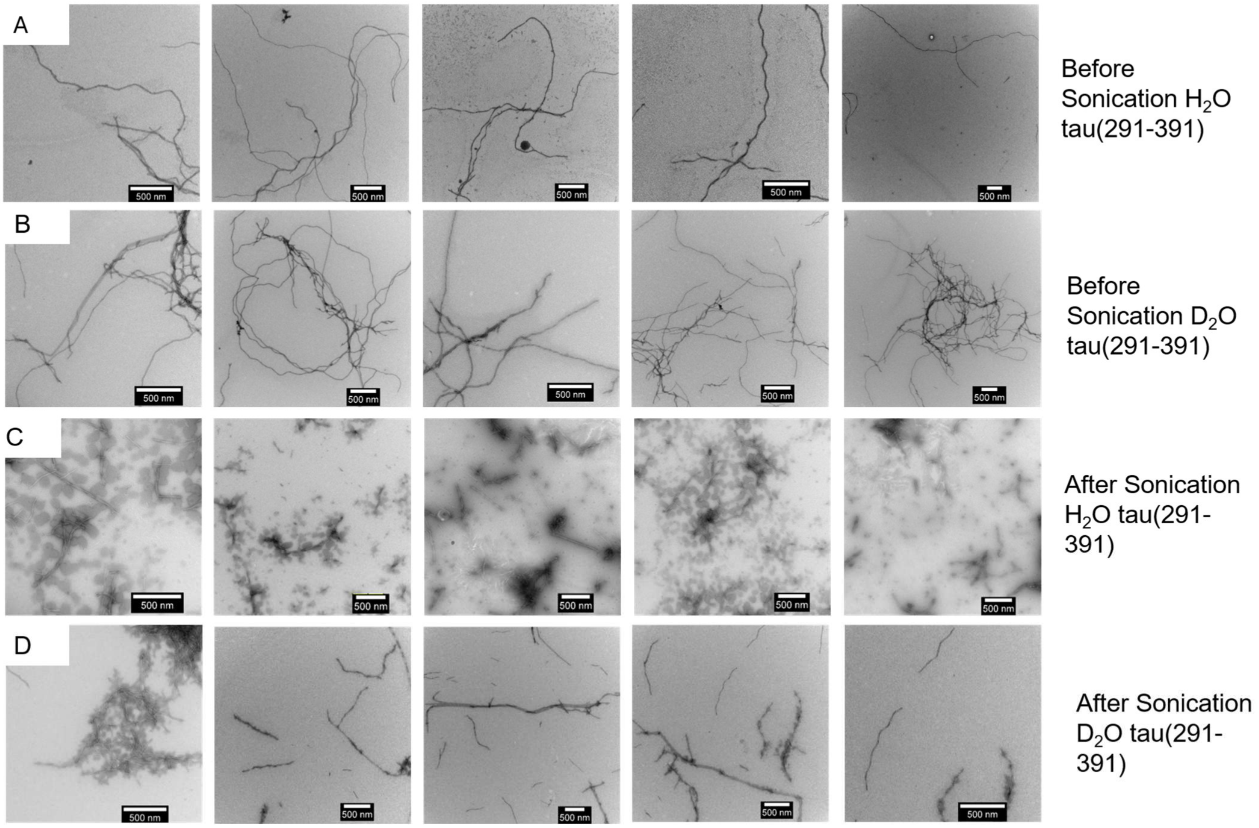

**Figure S3.** Solvent-dependent mechanical stability of tau(291–391) fibrils under sonication. Heparin-induced tau(291–391) fibrils were formed under $H_2O$ and $D_2O$ conditions (50 µM protein, 20 µM heparin, 10 mM DTT, 25 mM HEPES, pH/pD 7.4, 37 °C, 72 h). TEM images were acquired before (A and B) and after (C and D) sonication. Fibrils formed in $H_2O$ (C) display pronounced fragmentation into shorter species, whereas fibrils formed in $D_2O$ (D) retain longer and more intact structures. Scale bar: 500 nm.

### 8. Experimental Procedure for Proteinase K Digestion assay

An equal volume of fibrils of heparin induced tau(291-391) (50 µM monomers) and co-factor free tau(287-391)(6 mg/ml monomers) prepared in $D_2O$ or $H_2O$ was pelleted by centrifugation at 21000g for 30 min and probed with anti-DGAE antibody after denaturation in 8 M urea to detect the amount of tau fibrils in each solvent. Based on these results, for the proteinase K digestion assay, we performed fibril amount normalization by resuspending the pelleted fibrils in different amounts of 1x PBS to ensure that we treat an equal amount of fibrils from each solvent with Proteinase K. This is because fibril yields from the same concentration of monomers in different fibrillation reactions can be different.

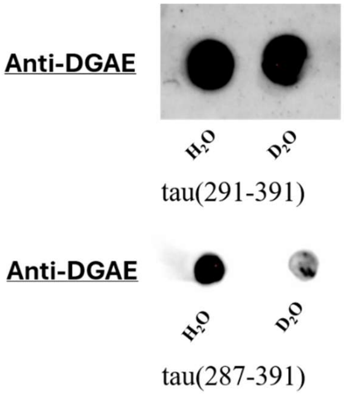

**Figure S4.** Anti-DGAE dot blot for plated heparin induced tau(291-391) and co-factor free tau(287-391) fibrils. Relative spot intensity was estimated using ImageJ for fibril dose normalization for $D_2O$ and $H_2O$.

This process also ensures that fibrils are freed from unrecruited monomers. Fibrils prepared in $H_2O$ or $D_2O$ were then incubated with Proteinase K at different ratios of fibril to Proteinase K at 37 °C for 30 min, and the digestion reactions were terminated using SDS-PAGE loading dye. SDS-PAGE was performed after heating the samples for 5 min at 90 °C.

#### 9. Quantification of Soluble Tau Fraction After Fibrilization

| Protein | $H_2O$:$D_2O$ ratio of soluble protein concentration in the supernatant after fibrillization |
|---|---|
| tau(287-391) | $H_2O$ vs $D_2O$ = 1.59:1 ± 4 |
| K18 | $H_2O$ vs $D_2O$ = 1.77:1 ± 8 |
| tau(291-391) | $H_2O$ vs $D_2O$ = 1.82:1± 3 |

**Table S1.** Quantification of soluble tau proteoforms remaining after aggregation in $H_2O$ and $D_2O$. Tau proteoforms were aggregated (50 µM protein, 20 µM heparin, 10 mM DTT, 25 mM HEPES (pH/pD 7.4), 37 °C. ) for 72 h followed by centrifugation, and the remaining soluble protein in the supernatant was quantified using a BCA assay. Values represent the relative amount of soluble protein remaining in $H_2O$ versus $D_2O$ from three independent experiments.

#### 10. Cell Seeding Experiment Details

HEK293T bioreporter cells expressing CFP/YFP K18 from ATTC () were used and maintained in Dulbecco's modified Eagle's medium (DMEM, high glucose) supplemented with GlutaMAX and 10% fetal bovine serum (FBS) kept at 37 in a humified incubator with 5% $CO_2$. 50,000 cells per well were transfected with 50 ul of mixture of proteoform stock. and lipofectamine 3000 in Opti-MEM. Transfection mixture was incubated at room temperature for 20 minutes prior to addition into cells for a final concentration of 50 nM of protein per well. Cells were incubated with the transfection mixture for 24 h at 37 °C. Cells were detached and resuspended in ice-cold 1x PBS. Single cell suspension was passed through 0.22 µm filter into flow cytometry tubes (BD FACSCelesta). FRET signals were acquired on a BD FACSCelesta flow cytometry. 10,000 events were collected for each technical replicate. Nine technical replicates per condition and three biological replicates were analyzed. Data was processed using FlowJo v10.

#### 11. Bio-Layer Interferometry

The bio-layer interferometry experiments were performed on a Sartorius Octet N1 system using Octet High Precision Streptavidin (SAX) biosensors. Sensors were hydrated in 25 mM HEPES (pH/pD 7.4) made either in $H_2O$ or $D_2O$. 4 µM heparin-biotin sodium salt conjugate (Sigma #B9806), or 5'-biotinylated U(40) (Genscript) was incubated for 45 minutes to immobilize the ligand. Following immobilization, the sensors were washed with HEPES for 2 minutes before being dipped into a concentration range of Tau. Binding kinetics were monitored and analyzed using the Octet system with a 5 min association/ 5 min dissociation time. Experiments were performed in triplicate with SD recorded. Protein dilution ranged from 20 µM – 0.3125 µM.

#### 12. Turbidity Experiment

Turbidity-based phase diagrams were obtained using a NanoDrop 1000 spectrophotometer. Absorbance was recorded at 350 nm using a path length of 0.1 cm. For each measurement, 3 µL of a 2x poly(U) RNA stock solution was combined with 3 µL of a 2x protein stock solution and gently mixed. After a 1 min equilibration period, 3 µL of the mixture was loaded onto the NanoDrop pedestal, and absorbance was measured twice. The same liquid–liquid phase separation (LLPS) mixture was measured once more to confirm reproducibility. Three independent biological replicates were performed for each condition. Absorbance values were averaged, and results are reported as SEM.

**Stock Solutions for R1-R8 either in $D_2O$ or $H_2O$:**
2x protein stock: -200 µM Proteins, and 25 mM HEPES (pH/pD 7.4).
2x poly(U) RNA stock: - 0, 20, 30. 60. 90, 120. 150, 180. 210. 240, 270. 300, 330, 360 µg/mL in 25 mM HEPES (pH/pD 7.4).

**Stock Solutions for Tau proteins either in $D_2O$ or $H_2O$.**
2x protein stock: -60 µM Tau, and 25 mM HEPES (pH /pD7.4).

**Final Concentration:**
1x Proteins, 1x µg/mL poly(U) RNA, 5 mM DTT, 25 mM HEPES (pH/pD 7.4).

**Turbidity experiment for R9-HBP-1**
Prepared a 6000 µM peptide stock of HBP-1 in 10 mM acetate buffer (pH/pD 4.5)
Mixed 1µL of peptide stock with 9 µL of 50 mM phosphate buffer (pH/pD 7.4) with varying salt concentrations (0, 0.4, 0.8, 1.0, 1.2, 1.6, 2.0 M NaCl) in 1:9 ratio.

**Final Concentration:**[4]
600 µM peptide, 0, 0.4, 0.8, 1.0, 1.2, 1.6, 2.0 M NaCl, 50 mM phosphate (pH/pD 7.4).

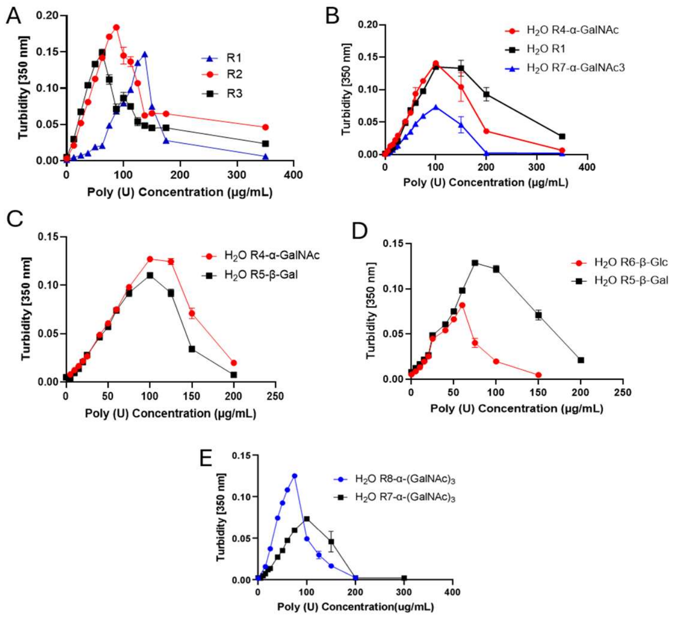

**Figure S5.** Comparison of turbidity among different RRASL peptide variants and glycopeptides at 100 µM peptide and variable concentrations of poly(U) in 25 mM HEPES buffer (pH/pD 7.4) at 23 °C in $H_2O$. (A) Comparison of R1, R2, and R3. (B) Comparison of R1, R4–α-GalNAc (mono-GalNAc), and R7–α-GalNAc$_3$ (tri-GalNAc). (C) Comparison of R4–α-GalNAc and R5–β-Gal. (D) Comparison of R5–β-Gal and R6–β-Glc. (E) Comparison of R7–α-GalNAc$_3$ and R8–α-GalNAc$_3$.

### 13. Circular Dichroism (CD) Spectroscopy

Proteins were dissolved in 20 mM sodium phosphate buffer containing 10 mM DTT (Ph/pD 7.4), prepared either in $H_2O$ or $D_2O$. Poly(U) was dissolved in 20 mM sodium phosphate buffer prepared in $H_2O$ or $D_2O$. Circular dichroism (CD) measurements were performed using a Modular Applied Photophysics Chirascan Plus CD spectrometer. Spectra were recorded in the far-UV region and averaged over two accumulations for each sample. A separate baseline was recorded under identical conditions and subtracted from the corresponding sample spectra.

The CD data were processed using Pro-Data Viewer software (Applied Photophysics) and converted to mean residue ellipticity (MRE). A smoothing factor of 10 was applied to improve the signal-to-noise ratio. The processed data were exported as text (txt) files using APL Data Converter and plotted using GraphPad Prism software.

**CD Spectrometer Settings**
Path length: 0.05 cm
Wavelength range: 180–280 nm
Step size: 0.5 nm
Bandwidth: 1 nm
Baseline: Separate baseline recorded for each measurement

### 14. Salt Resistance Assay of Proteins-RNA LLPS

The salt resistance assay was measured by turbidity on Nanodrop 1000. The absorbance was measured at 350 nm with a path length of 0.1 cm. A 0.65 µL Eppendorf tube was charged with 3 µL of 3x protein stock. The mixture was treated with 1.5 µL of 6x NaCl stock. Next, 4.5 µL of 2x RNA stock was added and pipetted up and down several times. A minute after mixing, 3 µL of the droplet solution was transferred to the Nanodrop, and the absorbance was measured three times. Using the same LLPS mixture, this was repeated twice more. This experiment was performed in triplicate, and the data from the three experiments was averaged and reported as SEM.

**Stock Solutions for R1:**
3x protein stock:
450 µM R1, and 25 mM HEPES (pH/pD 7.4).
6x NaCl stock: 150, 300, 450, 600 mM NaCl in 25 mM HEPES (pH/pD 7.4).
2x poly(U) RNA stock: 200 µg/mL poly U RNA in 25 mM HEPES (pH/pD 7.4).
**Final Concentration:**
- 100 µM R1, 100 µg/mL poly U RNA, 0-100 mM NaCl, 25 mM HEPES (Ph/pD 7.4), 23 ºC.
**Stock Solutions:**
3x protein stock:
-90 µM Tau Proteins, 15 mM DTT, and 25 mM HEPES (pH/pD 7.4). - 6x NaCl stock: 150, 300, 450, 600 mM NaCl in 25 mM HEPES (pH/pD 7.4). - 2x poly U RNA stock: 120 µg/mL poly U RNA in 25 mM HEPES (pH/pD 7.4).
**Final Concentration:**
- 30 µM K18, 60 µg/mL poly U RNA, 0-100 mM NaCl, 5 mM DTT, 25 mM HEPES (pH/pD 7.4), 23 ºC.

### 15. Labeling of tau(291-391) and K18

The labeling protocol is adapted from the AlexaFluor™ 488 5 N-Hydroxy Succinimidyl Ester (Thermo Scientific, Ref A20000) manufactures protocol. Protein solutions (6 mg/mL in $H_2O$) were adjusted to mildly basic conditions by the addition of $NaHCO_3$ (1 M, 3 µL) prior to reaction with AlexaFluor 488 dissolved in DMSO (10 mg/mL, 3 µL). The labeling reaction was allowed to proceed for 2 h at room temperature (23 °C) and was subsequently terminated by the addition of hydroxylamine (50% w/v, 10 µL). Unreacted fluorophore was removed by size-exclusion desalting on Sephadex G-25 equilibrated with $H_2O$. Fractions containing protein were identified by gel electrophoresis, pooled, and lyophilized. The dried material was reconstituted in 25 mM HEPES and subjected to further purification by dialysis against 25 mM HEPES (pH/pD 7.4) using a 2 kDa molecular weight cutoff Slide-A-Lyzer™ MINI dialysis device. Following 48 h of dialysis, protein concentrations were determined using a BCA assay, while fluorophore concentrations were quantified by UV–vis spectroscopy at 495 nm ($\varepsilon$ = 71,000 $M^{-1}$ $cm^{-1}$). The degree of labeling was calculated from absorbance measurements and was approximately one fluorophore per protein molecule for both Tau (291–391) and K18 constructs see table below.

| Proteins | MW [g/mol] | Tau (BCA) [µM] | AF488 (A495) [µM] |
|---|---|---|---|
| tau(291-391) | 10770.2 | 125 | 141 |
| K18 | 13723 | 22 | 18 |

### 16. FRAP of Tau and RRASL (R1-R7) Constructs

For FRAP measurements, 3 µL of a 2x protein stock prepared in either $H_2O$ or $D_2O$ was mixed with 3 µL of a 2x poly(U) RNA stock prepared in the corresponding solvent. The solutions were gently mixed by repeated pipetting and immediately transferred to the center of a micro-well in a 35 mm glass-bottom dish containing a 14 mm well. To minimize evaporation, approximately 12 µL of 25 mM HEPES buffer was placed around the perimeter of the micro-well. The chamber was covered with a glass coverslip and sealed using nail polish. Samples were allowed to equilibrate for 1 h prior to photobleaching. For each droplet preparation, a single condensate was bleached and monitored for fluorescence recovery. All measurements were performed in three independent experiments.

**Stock Solutions of Tau:**
2x protein stock: 60 µM tau(291-391) and K18, 0.6 µM AF488, 20 mM DTT, and 25 mM HEPES (pH/pD 7.4).
2x poly(U) RNA stock: 80 µg/mL poly(U) RNA in $H_2O$ and 60 µg/mL in $D_2O$ for Tau 291–391; for K18, 120 µg/mL poly(U) RNA in $H_2O$ and 90 µg/mL in $D_2O$, all prepared in 25 mM HEPES (pH/pD 7.4).
**Final Concentration:**
30 µM tau(291-391) and K18, 0.3 µM AF488 (labelled tau(291-391)), 1x µg/mL poly(U) RNA, 10 mM DTT, 25 mM HEPES (pH/pD 7.4), 23 ºC.
**Stock Solutions of R1:**
-2x protein stock: 300 µM of R1, 30 µM R1-FITC-labelled, and 25 mM HEPES (pH/pD 7.4).
-2x poly(U) RNA stock: 300 µg/mL poly(U) RNA either in $H_2O$ or $D_2O$ all prepared in 25 mM HEPES (pH/pD 7.4).
**Final Concentration:**
150 µM of R1 0.15 µM R1-FITC-labelled 150 µg/mL poly(U) RNA, 25 mM HEPES (pH/pD 7.4), 23 ºC either in $H_2O$ or $D_2O$.

**Stock Solutions of R5, R6, and R7:** 2x protein stock: 300 µM of R5, R6, and R7, 20 % v/v FITC-labelled RNA + 300 µg/mL poly(U) RNA, and 25 mM HEPES (pH/pD 7.4) either in $H_2O$ or $D_2O$, 20 % v/v FITC-labelled RNA always in $H_2O$.

**Final Concentration:** 150 µM of R5, R6, and R7 10 % v/v FITC-labelled RNA + 150 µg/mL poly(U) RNA, and 25 mM HEPES (pH/pD 7.4) ,10 % v/v FITC-labelled RNA always in $H_2O$, 23 ºC either in $H_2O$ or $D_2O$.

**FRAP Microscope Settings**
FRAP experiments were conducted using a Nikon Eclipse Ti laser scanning confocal microscope equipped with a 60× oil-immersion objective. Condensates were visualized in the EGFP channel using a laser power setting of 3 and a detector gain of 5, with the pinhole set to 1.2. Images were acquired at Nyquist sampling (0.09 µm pixel size) with a 1024 × 1024 frame. All droplets were imaged at the bottom surface of the dish using the Perfect Focus System (PFS), and acquisition settings were automatically scaled to prevent signal saturation.

Photobleaching was performed using the 488 nm laser line at a power setting of 20. Circular regions of interest (ROIs) were defined for stimulation, background, and reference measurements, each with identical Feret dimensions (1.2 × 1.2). Fluorescence recovery data were processed in GraphPad Prism using a double-normalization procedure as described in the literature.[5] Data from three independent replicates were averaged and plotted, with error bars representing as SEM.

**Mobile Fraction Calculation**
The mobile fraction (MF) was calculated according to the following equation:

$$MF = \frac{I_\infty - I_c}{I_{c_0} - I_c}$$

where $I_\infty$represents the fluorescence intensity at the recovery plateau, $I_c$is the fluorescence intensity immediately after photobleaching, and $I_{c_0}$is the fluorescence intensity prior to photobleaching. Mobile fractions are reported as percentages.

**Timing for FRAP experiments for Tau:**
- Acquisition after 2 second intervals over 10 seconds (5 data points).
- Stimulation for 2 seconds.
- Acquisition after 2 second intervals over 1 minute 30 seconds (45 data points).
- Acquisition after 10 second intervals over 7 minutes (42 data points).

**Timing for FRAP experiments for R1, R5, R6, and R7:**
- Acquisition after 2 second intervals over 4 seconds (2 data points).
- Stimulation for 2 seconds.
- Acquisition after 2 second intervals over 1 minute (25 data points).

### 17. Droplet Size Measurements

Droplet size measurements were performed using a Nikon Eclipse Ti laser scanning confocal microscope equipped with a 60× oil-immersion objective. Droplets were imaged in the EGFP channel using a laser power setting of 3 and a detector gain of 5. The pinhole size was set to 1.2. All imaging fields were acquired at Nyquist sampling (0.09 µm pixel size) with a 1024 × 1024 resolution.

Images were analyzed using ImageJ software following this workflow: Open Image → Convert to 8-bit → Threshold → Huang threshold → Analyze Particles (size = 0.1–∞, circularity = 0–1; overlay masks displayed; results summarized; edge particles included). The resulting average droplet area values were plotted graphically, with error bars representing the SEM.
